# Functional Category-Specific Intolerance Reflects Genic Function and Clinical Relevance

**DOI:** 10.1101/2025.09.24.678298

**Authors:** Yuncheng Duan, Constantine Stavrianidis, Grace Tzun-Wen Shaw, Christopher Sottolano, Ramakrishnan Rajagopalan, Tanaya Jadhav, Grace E. Rhodes, Tristan J. Hayeck, Andrew S. Allen

## Abstract

A key problem in genetics is associating variants with disease phenotypes. In aid of this, much progress has been made in quantifying the functional impact of individual variants on the gene product it codes for. However, the intolerance of the sequence in which those variants are found to functional variation is also a key determinant of whether a deleterious variant is pathogenic or not. Previous approaches to estimating genic intolerance have combined functional variant types, i.e., missense, loss-of-function, etc., or restricted analyses to only one type, i.e., pLI, missense-Z etc. Here we take a different approach and jointly model patterns of intolerance across multiple functional variant types. We refer to this approach as CATMINT. We show that CATMINT is competitive with previous gene level intolerance metrics in predicting disease relevant genes, with CATMINT ranking among the top performing scores across differing types of genes. However, perhaps more exciting is that CATMINT enables variant category specific intolerance estimation, revealing distinct functional profiles across genes/gene families. Analysis of ClinVar data shows that CATMINT intolerance patterns in disease genes recapitulate patterns of pathogenic variants within those genes, supporting the utility of category-specific intolerance in clinical variant interpretation. Further, we use the statistical framework utilized by CATMINT to conduct power analyses, allowing us to classify genes according to the power those genes have to detect intolerance. This allows us, for example, to identify genes that are underpowered and undetected, but may nevertheless be highly intolerant. Together, these results define a framework for understanding how selective pressures shape gene-specific sensitivity to different classes of mutation, improving the resolution of variant interpretation and gene prioritization in clinical and functional genomics.

## Introduction

Identifying genetic coding variation under purifying selection has important implications for understanding disease mechanisms and improving genetic diagnoses. For example, knowing that a certain type of variant is selected against in a given region of a Mendelian disease gene may help with diagnosis when such a variant is observed in a patient presenting with genetic disease. Signals of selection can be detected at different evolutionary scales, including both inter-species conservation and intra-species intolerance or constraint. Cross species conservation, i.e., the extent to which a sequence is more similar across species than expected by chance, has been widely used to identify functionally important regions. Conservation can be evaluated through comparisons of orthologous sequences across species or by measuring deviations of observed substitutions from expectation across phylogenies (Christmas et al., 2023; Cooper et al., 2005; Davydov et al., 2010; Genereux et al., 2020; Siepel et al., 2005).Click or tap here to enter text.

However, conservation-based approaches are relatively insensitive to recent, usually species specific, selection. An alternative approach is to use genetic intolerance or constraint, i.e., the depletion of variation relative to expectation in a large cohort of individuals from the same species (Hayeck et al., 2019; Karczewski et al., 2020; Lek et al., 2016; Petrovski et al., 2013). Intolerance has at least two advantages over conservation. First, it can capture sequences that have gained functional importance more recently, including those unique to a species or a small phylogenetic branch. Second, its power scales with the size of the cohort. With the increasingly large human cohorts available today, intolerance approaches can identify regions subject to far more subtle selection effects. For example, there are 730,947 whole exomes and 76,215 whole-genomes from unrelated individuals across multiple populations in gnomAD V4 (Karczewski et al., 2020).

Multiple intolerance approaches have been developed to estimate the degree of purifying selection based on variation observed within human populations. One class of methods, referred to here as “conditional” approaches (Bustamante et al., 2005; Gussow et al., 2016; Hayeck et al., 2019; Petrovski et al., 2013), infers selection by comparing the number of observed functional variants to the total amount of variation in a gene. For example, RVIS (Petrovski et al., 2013) uses a linear regression framework to evaluate the deviation of common functional variants per gene from genome-wide average, given the gene’s total variation. This framework was further extended to subregions of genes, such as in subRVIS and LIMBR (Gussow et al., 2016; Hayeck et al., 2019). A limitation of conditional approaches is that they do not explicitly account for the site-specific mutability, which can vary due to the local genetic context. In contrast, a different approach, which we call the “mutability” approach, estimates the expected number of variants in a certain variant class based on the sequence context mutability of the class. For example, trinucleotide sequence context mutability has been used to model depletion of missense variants (missenseZ, measured by the gnomAD group, and in Samocha et al., 2014), protein truncating variants (pLI, pRec, pNull; Lek et al., 2016) and loss-of-function variants – pLOF and LOEUF (Samocha et al., 2014; Karczewski et al., 2020).

Because mutability approaches estimate expectations using sequence-context mutability models, their computations are restricted to very rare variation where demographic biases are minimized. As a result, only part of the genetic variation is used, with common variants being disregarded, negatively impacting power. In contrast, in conditional approaches, the number of observed functional variants are compared to genome-wide average conditional on the total number of observed variants per gene. This comparison is less susceptible to demographic bias, since demographic factors would impact both functional and nonfunctional variation similarly. As a result, conditional approaches can incorporate a broader spectrum of observed variants into the analysis, including common ones, which could improve power.

In this study, we integrate the strength of the two approaches by incorporating sequence-context mutability into a conditional modeling framework. In addition, we also incorporate functional information that can further enrich the constraint/intolerance measurements by infusing biologically meaningful information into the modeling to assess intolerance across functional categories. We classified variants into five distinct functional categories, i.e., codon-optimality-neutral (CON), codon-optimality-change (COC), less-damaging-missense (LDM), probably-damaging-missense (PDM), and loss-of-function (LOF), and jointly analyzed their intolerance. This classification reflects the varying impacts of different variant types on gene function (Gerasimavicius et al., 2022) and their relevance to human diseases. By conducting inference across variant types jointly within the same statistical model, we can characterize the global shift in variant composition relative to neutral expectation and generate more precise and informative genic intolerance estimates. More importantly, because the joint model provides estimates for each category, we can compare their relative depletion and formally test for differences. This allows us to conduct direct inference on how natural selection shapes the genetic architecture of each gene based on their functional roles.

## Results

### Improved Gene Overall Intolerance Assessment with LRT-Based Modeling in CATMINT

To generate gene-level intolerance scores based on the joint estimation across multiple variant categories, we performed CATMINT (CATegory-based Multinomial likelihood ratio test for INTolerance, Method) on genes that are included in the Consensus Coding Sequence (CCDS) sets (Pujar et al., 2018). Specifically, we classified all potential single nucleotide changes from reference — three per base within each transcript — into the following categories, based on their functional consequence: codon optimality neutral (CON), codon optimality change (COC), less damaging missense (LDM), probably damaging missense (PDM), and loss-of-function (LOF) (Method). Given the categorical nature of these classifications, the data inherently aligns with analysis using a multinomial model (Figure 1A).

**Figure 1.**
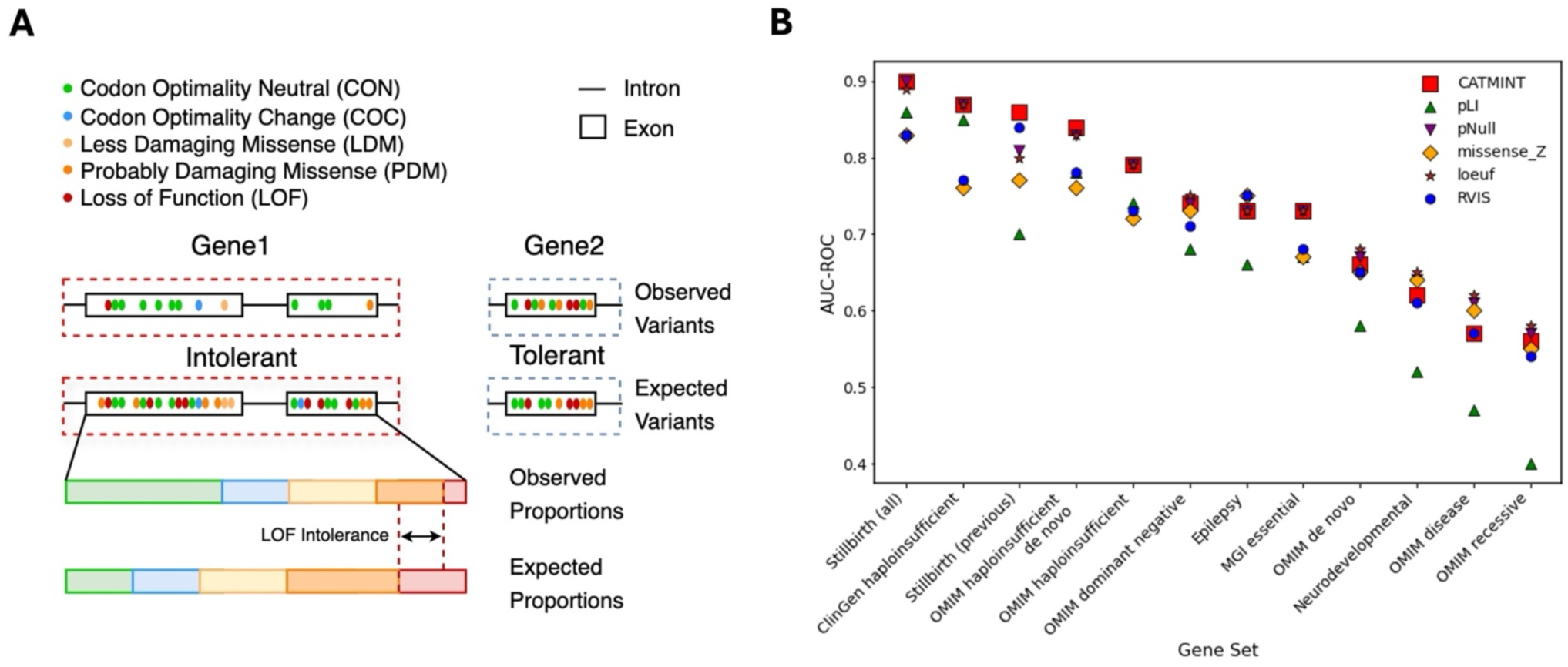
CATMINT Framework and Performance Benchmarking Against Established Gene Intolerance Metrics. **(A)** Schematic of the CATMINT framework. CATMINT models gene-level intolerance by comparing the observed composition of variants (from gnomAD v4) to expected proportions based on sequence-context–dependent mutability under neutrality. Variants are classified into five functional categories: codon optimality neutral (CON), codon optimality change (COC), less damaging missense (LDM), probably damaging missense (PDM), and loss-of-function (LOF). The schematic shows an intolerant gene (Gene1) with a depletion of proportion in functional variants, e.g., LOF variants, relative to expectation, and a tolerant gene (Gene2) with composition more closely matching the neutral expectation. Deviations in variant composition form the basis of the likelihood ratio test. **(B)** Comparison of CATMINT to established intolerance scores across 12 curated gene sets. Gene sets include those from OMIM (e.g., haploinsufficient, de novo, dominant negative), ClinGen haploinsufficient genes, neurodevelopmental and epilepsy genes, MGI essential genes, and stillbirth-associated genes. CATMINT (red square) consistently ranks among the top-performing methods in most gene sets compared to other metrics — including pLI, pNull, missense_Z, loeuf, and RVIS, particularly those under strong purifying selection (e.g., stillbirth and haploinsufficient genes). AUC values across methods and gene sets could be found in Supplementary Table 6. *In addition to stillbirth (all) genes, we also generated stillbirth (previous) genes: excluding the stillbirth candidate genes, to prevent bias from enrichment of candidate genes with lower LOEUF scores (Method).

CATMINT evaluated the deviation in a gene’s standing variant composition from neutral expectation, based on the relative mutability of each variant type. The mutability, or the neutral variant rate, was estimated empirically from putative neutral sequence (Method). Standing variants were defined as gnomAD v4 (Karczewski et al., 2020) exome variants meeting quality control criteria, representing variation in the healthy human population (Method). The expected number of variants in each category was calculated using the sequence context dependent variant rate (Method) under neutrality.

We treated CATMINT-derived p-values as gene-level intolerance scores (CATMINT measurements for all protein-coding genes could be found at Supplementary Data 1). Validation of these scores was conducted against various gene sets from the Online Mendelian Inheritance in Man (OMIM) database (Amberger et al., 2009), extracted using keywords search, following the instruction in Petrovsky et al., 2013. Additional gene sets are included such as epilepsy, neural developmental (Hayeck et al., 2019), still birth (Stanley & Goldstein, 2020), ClinGen haploinsufficient genes (Rehm et al., 2015), and MGI (Mouse Genome Informatics) essential genes (Blake et al., 2011; Georgi et al., 2013; Liu et al., 2013). A control set was also defined, excluding genes from all other sets (Method).

We generated Receiver Operating Characteristic (ROC) curves for those gene sets and computed the area under the receiver operating curve (AUC) of each gene set (Supplementary section 5; Supplementary Figure 4), demonstrating that CATMINT scores could distinguish each gene set from the control with varying degrees of accuracy. In general, gene sets expected to be under strong purifying selection yielded higher AUC values, such as stillbirth genes (AUC=0.90 [95% CI: 0.86-0.95]). Genes associated with fetal death should be under strong purifying selection since stillbirth causing mutations are likely to greatly impact fitness. Similarly, the intolerance scores exhibited high accuracy in discriminating haploinsufficient (HI) gene sets from the control set, with ClinGen HI genes showing an AUC of 0.87 (95% CI: 0.84-0.90), and an AUC of 0.84 (95% CI: 0.81-0.86) for OMIM haploinsufficient de novo genes. Conversely, the olfactory receptor genes, known to be highly polymorphic among humans and other species, show less deviation from neutrality in variant composition compared to the control gene set (AUC = 0.25 [95% CI: 0.23-0.27]).

We benchmarked CATMINT against other known gene intolerance scores such as RVIS (Petrovski et al., 2013), pLI, pNull, missense_Z, and loeuf (Karczewski et al., 2020), using the area under the receiver operating characteristic (AUC-ROC) curves for each metric. To ensure comparability across different scores, we recalculated the RVIS score using the gnomAD V4 whole exome data, updating from the original 2012 dataset (NHLBI Exome Sequencing Project (ESP); (Fu et al., 2013; Tennessen et al., 2012)) which was significantly smaller. This recalculation and all quality control measures were performed in accordance with the procedures we established for CATMINT (Supplementary Section 6). These analyses revealed that CATMINT ranked consistently among the top performers across all evaluated gene sets and outperformed the established scores in the majority of them (Figure 1B, Supplementary Table 6). These results support the robustness of our model and its utility in quantifying gene intolerance to functional variation.

We calculated the enrichment fold of the top intolerant genes within each gene set, with the top intolerant genes defined as those falling within percentiles ranging from the top 1% to the top 50% based on CATMINT p-values (Supplementary Section 5). ClinGen HI genes, OMIM HI de novo genes, and stillbirth genes show a dramatic enrichment with over 15-fold in the top 1% intolerant genes. Similarly, OMIM HI, and epilepsy genes are also significantly enriched, with more than 10-fold enrichment in the top 1% intolerant genes, while recessive genes slightly depleted in top 1% intolerant genes (Supplementary Figure 5).

### Variant Type Specific Intolerance Uncovers Functional Mechanisms of Genes

CATMINT not only provides accurate prediction of the overall functional intolerance but also evaluates intolerance across multiple variant categories simultaneously. To quantify category-specific intolerance, we fit a baseline category logit regression model with codon optimality neutral (CON) variants as the baseline category and calculated the log-odds (*δ*) and its z-score for each functional variant type (Method). A positive *δ* or z-score indicates that the variant category is relatively more tolerant than CON variants, suggesting possible tolerance to that category. Conversely, a negative *δ* or z-score indicates intolerance in this variant category relative to CON variants, with more negative values reflecting stronger intolerance and potential gene functions disrupted by such variants.

To test whether patterns of category-specific intolerance reflect gene function, we evaluated the enrichment of category-specific intolerant genes across super gene families (SGFs). For each variant category, we labeled genes as intolerant which have a z-score in the lowest 10^th^ percentile. Genes were grouped according to their SGFs, and enrichment of intolerant genes was computed (Method).

A higher enrichment score of a SGF for a given variant category indicates that genes in that family are particularly intolerant to mutations of that type. While we’ve calculated enrichment of intolerant genes for each SGF (Supplementary Data 2), here for illustration, Table 1 highlights selected SGFs with distinct intolerance profiles, including COC-specific, LOF-specific, PDM-specific, and PDM/LOF-specific intolerance (Method). The variant intolerance patterns reflect the functional distinctions across gene families.

**Table 1.**
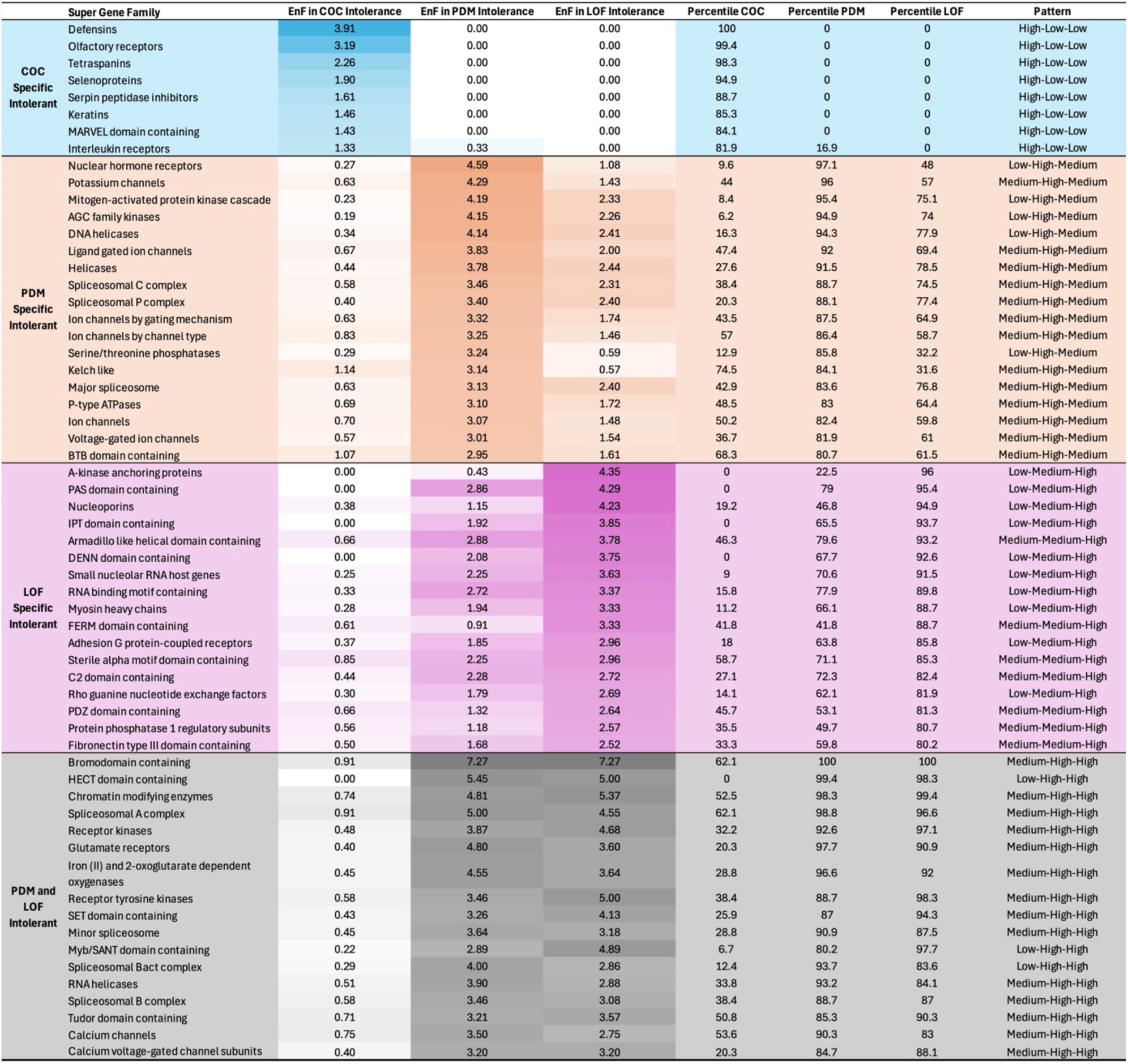
Distinct intolerance patterns across Super Gene Families (SGFs). SGFs were ranked by EnF (enrichment fold) in COC, PDM, and LOF variants, respectively. High/Medium/Low labels for each variant category were assigned by quantile rank threshold (>80th percentile: High; <20th percentile: Low; Otherwise: Medium), and representative SGFs were selected based on the specific label patterns (COC specific: H-L-L; PDM specific: L-H-M, M-H-M; LOF specific: L-M-H, M-M-H; PDM/LOF specific: L-H-H, M-H-H). Within each variant specific intolerance group, SGFs were ranked by enrichment in the defining variant category (PDM/LOF specific SGFs were ranked by the average of enrichment of PDM and LOF). The EnF values for each variant category are color-coded within each intolerance group, with darker shades indicating stronger enrichment.

**Table 2.**
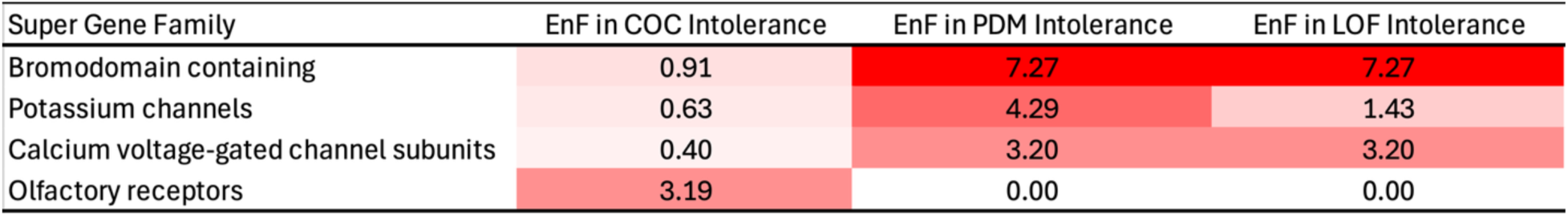
Selected Super Gene Families (SGFs) with distinct intolerance patterns. Shown here are selected SGFs extracted from Table 1, including bromodomain-containing genes and calcium voltage-gated channel subunit genes (intolerant to both PDM and LOF), potassium channel genes (intolerant to PDM), and olfactory receptor genes (intolerant to COC).

| Super Gene Family | EnF in COC Intolerance | EnF in PDM Intolerance | EnF in LOF Intolerance |
| --- | --- | --- | --- |
| Bromodomain containing | 0.91 | 7.27 | 7.27 |
| Potassium channels | 0.63 | 4.29 | 1.43 |
| Calcium voltage-gated channel subunits | 0.40 | 3.20 | 3.20 |
| Olfactory receptors | 3.19 | 0.00 | 0.00 |

SGFs with strong intolerance to COC while showing little to no intolerance to PDM or LOF variants were categorized as COC-specific intolerant SGFs. This pattern suggests that these genes might be particularly sensitive to translational efficiency, mRNA stability, or other post-transcriptional regulations, rather than to disruptions in protein sequence or complete gene loss, potentially due to tight expression control or specialized translational mechanism. Examples SGFs include olfactory receptor genes and defensins. PDM-specific intolerant SGFs often include genes encoding proteins with significant structural roles, such as ion channels, kinases, or spliceosome components, where amino acid substitutions can disrupt their functions. In contrast, LOF-specific intolerant SGFs include genes with dosage-sensitive roles. They are enriched for signaling and regulatory roles, where gene loss is relatively more deleterious than missense (mostly structural) changes. Notably, LOF and PDM intolerance are not independent: no SGF shows high LOF intolerance without at least moderate PDM intolerance, and vice versa. Thus, the terms “LOF-specific” or “PDM-specific” reflect relative intolerance. Last, SGFs that showed strong intolerance in both PDM and LOF variants indicate sensitivity in both protein structure and dosage. These families include genes involved in epigenetic regulation, development, and core cellular processes.

### ClinVar Variant Analysis Confirms the Diagnostic Application of Category-Specific Intolerance in PDM and LOF

Variant category specific intolerance can improve variant interpretation and candidate gene prioritization in clinical diagnostics. For example, if a gene is more intolerant to probably damaging missense (PDM) variants than to loss-of-function (LOF) variants, PDM mutations in that gene are more likely to be pathogenic, and vice versa for genes showing stronger LOF intolerance. In clinical exome sequencing, category-specific intolerance can help prioritize candidate genes by linking observed variant types to a gene’s specific intolerance profile. This approach moves beyond the focus on LOF variants. For example, if a patient presents with an enrichment of PDM or codon-optimality-change (COC) variants in a gene known to be intolerant to those types, it provides additional support for pathogenicity even in the absence of LOF mutations.

For each gene in the high confidence set, we measured the relative intolerance between PDM and LOF variants using a contrast derived from the category logit model presented earlier (Method). To evaluate whether this contrast reflects clinically relevant patterns, we extracted ClinVar (Landrum et al., 2014) pathogenic and likely-pathogenic variants and counted the number of PDM and LOF variants per gene.

Genes with more ClinVar pathogenic missense variants tended to show stronger intolerance to PDM variants from the gnomAD population (Figure 2A). In contrast, no such trend was observed for ClinVar variants with uncertain significance or classified as benign or likely benign (Supplementary Figure 6), which is consistent with the expectation that these variants are not under strong selective pressure.

**Figure 2.**
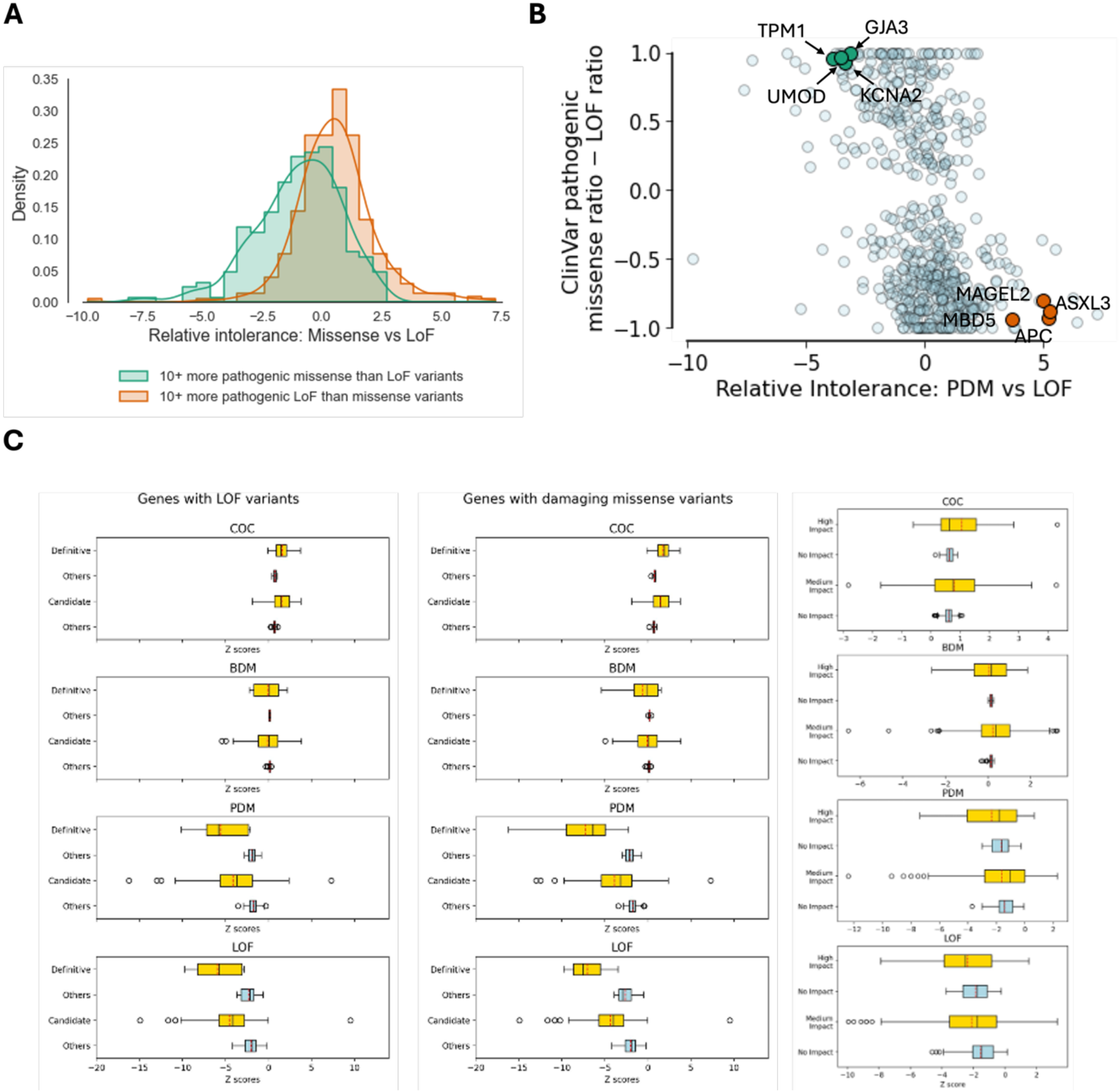
Variant-category-specific intolerance informs variant interpretation and clinical prioritization. **(A)** Genes grouped by ClinVar pathogenic variant imbalance show distinct patterns of relative intolerance. Genes with ≥10 more pathogenic missense than LOF variants (green) and those with ≥10 more pathogenic LOF than missense variants (orange) are compared using their CATMINT-derived contrast scores (*z*_PDM−LOF_). Genes enriched for missense pathogenic variants show stronger relative missense intolerance, while those enriched for LOF variants tend to show stronger LOF intolerance. **(B)** Correlation between CATMINT contrast scores (*z*_PDM−LOF_)and the difference in proportions of ClinVar pathogenic missense and LOF variants per gene. The x-axis shows the difference in ClinVar pathogenic missense vs. LOF variant proportions, and the y-axis shows the relative intolerance (*Z*_PDM−LOF_). Genes with higher pathogenic missense burdens tend to show stronger missense intolerance, while genes with higher LOF burdens show stronger LOF intolerance (Pearson’s r = –0.44, p < 2.2×10⁻¹⁶). Highlighted examples include missense-intolerant genes (e.g., TPM1, UMOD, KCNA2, GJA3) and LOF-intolerant genes (e.g., MBD5, MAGEL2, ASXL3, APC). **(C)** Intolerance scores for CHD-associated gene sets across variant impact categories. Left: Box plots comparing intolerance z-scores across four CATMINT-based annotation tests for known loss-of-function (LOF) CHD-associated genes. Within each annotation, distributions are shown separately for definitive genes (n=13) and candidate genes (n=104). Middle: Similar analysis for CHD-associated missense genes (14 definitive, 103 candidate), showing intolerance across the same CATMINT annotation schemes. Right: Intolerance z-score distributions for genes harboring high-impact or moderate-impact de novo variants in the BDB cohort. Genes in panel are compared to a length matched permutated set of genes.

We further quantified this relationship by comparing the difference in intolerance between PDM and LOF variants (*Z*_!“#$%&’_) with the corresponding difference in the proportions of ClinVar pathogenic variants across the two categories. The two measures showed a moderate but significant negative correlation (Figure 2B). Pearson’s cor = −0.44, CI: −0.50 – - 0.37, p-value < 2.2e-16), indicating that stronger relative intolerance to PDM is associated with a higher proportion of ClinVar pathogenic missense variants.

Missense-intolerant genes, such as TPM1, UMOD, GJA3, and KCNA2, showed strong depletion of missense variants relative to LOF variants and carried a disproportionately high number of pathogenic missense variants in ClinVar (Figure 2B). These genes often encode proteins with structurally or mechanistically sensitive domains, such as channel genes, where amino acid substitutions can perturb function. For instance, pathogenic missense mutations in TPM1, which encodes cardiac tropomyosin, impair thin-filament regulation and actin-myosin interaction, leading to inherited cardiomyopathies (Olson et al., 2001). In GJA3, missense variants disrupt gap-junction channels in the lens and cause congenital cataracts (Goyal et al., 2024). Similarly, missense variants in KCNA2, a potassium channel gene, are known to alter gating properties, causing epileptic encephalopathy, through a dominant negative effect (Masnada et al., 2017; Syrbe et al., 2015). In UMOD, missense variants disrupt protein folding and cause accumulation of misfolded uromodulin in the endoplasmic reticulum, leading to autosomal dominant tubulointerstitial kidney disease (Lens et al., 2005; Rampoldi et al., 2003).

Conversely, genes such as MAGEL2, ASXL3, MBD5, and APC showed stronger depletion of LOF variants and were enriched for pathogenic LOF mutations in ClinVar (Figure 2B). Many of those genes are known to be dosage-sensitive and associated with developmental disorders or cancer predisposition and tend to cause disease through haploinsufficiency. For example, MAGEL2 is a paternally expressed, imprinted gene where LOF mutations cause Schaaf–Yang syndrome, with features including developmental delay and autism spectrum traits (Huang et al., 2023). Its monoallelic expression likely amplifies its LOF intolerance. ASXL3, a chromatin remodeling factor, is disrupted in Bainbridge–Ropers syndrome, where truncating variants lead to severe neurodevelopmental impairment (Woods et al., 2024). Both genes show strong depletion of LOF variation in the population and an enrichment of pathogenic LOF variants in ClinVar, consistent with their clinical impact. These patterns demonstrate that category-specific intolerance can help prioritize disease-causing variants by linking observed variant types to gene-specific sensitivity.

Relative intolerance also agrees well with a ClinVar-based strategy for drug discovery. Ressler and Goldstein (Ressler & Goldstein, 2023) used a high ClinVar missense ratio (>0.8) and low pLI (<0.1) to highlight 36 autosomal dominant genes likely affected by gain-of-function mechanisms. These genes generally show more negative *Z*_*PDM*−*L*0*F*_ in our model, indicating greater PDM than LOF intolerance (Supplementary Data 3). In contrast, genes with high ClinVar missense ratio (>0.8) while having high pLI (>0.1; 85 genes) show intolerance across both categories (Supplementary Figure 7; Supplementary Data 3). This concordance shows that relative intolerance captures similar signals to the ClinVar ratio, while offering potential advantages in settings where ClinVar is biased toward well-studied genes or limited by sparse variant counts, or the inheritance mode is unclear. Some additional genes, such as TPM1, also show patterns consistent with gain-of-function potential but were not included in the 36 genes identified by Ressler and Goldstein. Z-scores for PDM, LOF, and relative intolerance are provided for all protein-coding genes (Supplementary Data 1) and can be used to evaluate potential mechanisms of genes of interest. All three scores should be considered together to distinguish gain-from loss-of-function mechanisms.

### De Novo Variants Associated with CHD Are More Intolerant in PDM and LOF

To investigate the potential impact of our intolerance scores on disease gene mapping, we analyzed known CHD-associated (Morton et al., 2022) or enriched genes with either LOF variants or missense variants. These gene sets were further subdivided into definitive (statistically significant) and candidate (enriched in variation but not statistically significant:13 definitive LOF genes and 14 definitive missense genes, 104 candidate LOF genes and 103 candidate missense genes. Paired t-tests were performed to compare CHD associated genes to non-CHD genes matched on coding sequence (CDS) length. Standard t-tests were performed when comparing definitive versus candidate genes since matching on gene length is problematic in this setting (Figure 2C).

LOF genes showed significantly more intolerance in both LOF (definitive: mean difference = –3.6, p-value = 5.3 × 10⁻⁵; candidate: mean difference = –2.5, p-value = 8.5 × 10⁻¹⁸) and PDM (definitive: mean difference = –3.6, p-value = 9.2 × 10⁻⁵; candidate: mean difference = –2.3, p-value = 1.3 × 10⁻¹¹) compared to length-matched non-CHD genes. Without matching on gene length (i.e. using unpaired t-test), definitive genes were significantly more intolerant to LOF than candidate genes (mean difference = −1.4, p-value = 0.05).

CHD-associated missense genes were also significantly more intolerant than length-matched non-CHD genes in both PDM (definitive: mean difference = 5.2, p-value = 5.3 × 10⁻⁵; candidate: mean difference = 2.1, p-value = 4.1 × 10⁻¹¹) and LOF (definitive: mean difference = 4.3, p-value = 2.5 × 10⁻⁷; candidate: mean difference = 2.4, p-value = 3.0 × 10⁻¹⁶). Definitive genes were significantly more intolerant than candidate genes for both PDM (mean difference = 4.3, p-value = 8.5 × 10⁻⁵) and LOF (mean difference = 2.7, p-value = 4.7 × 10⁻⁴) based on their z-scores.

We also analyzed a second gene set derived from de novo variants identified in CHD probands from 533 high-coverage sequencing proband-parent trios from the Birth Defects Biorepository (BDB) at the Children’s Hospital of Philadelphia. We grouped genes based on their de novo variant’s highest impact. Genes with at least one moderate-impact variant are significantly more intolerant to LOF (mean difference = −0.56, p-value = 2.6 × 10⁻⁵) compared to the no-impact, coding-domain sequence length-matched genes. Trends observed in the BDB gene sets were generally similar to the published CHD genes: the high-impact distributions resembled those of the definitive CHD genes, showed the strongest intolerance while the moderate-impact intolerance was similar to those of the CHD candidate genes, as seen in both PDM z-scores and LOF z-scores.

### Depletion in Codon Optimality Change Variants Indicates Limited Synonymous Intolerance in Specific Gene Sets

Variants that alter codon optimality can affect mRNA stability, protein translation efficiency and other critical processes (Presnyak et al., 2015). Previous studies have suggested that certain gene sets might be particularly sensitive to these types of variants, such as DNA repair genes, genes associated with G1/S cell replication phase, and dosage sensitive genes (Dhindsa et al., 2020). To assess this, we used z-scores of codon optimality change (COC) variants derived from our baseline category logit model presented above to measure the depletion of COC variants in comparison to the codon optimality neutral (CON) variants.

None of the three gene sets showed significant intolerance to COC variants, as reflected in both the z-score distributions (Figure 3A) and AUCs (Supplementary Figure 8). These results do not align with previous findings (Dhindsa et al., 2020), which reported stronger intolerance using synRVIS, a score that measured the depletion in the optimal to non-optimal (OP -> NO) variant given total synonymous variants in a gene. In our model, COC category includes both directions of codon optimality change (OP -> NO and NO -> OP). To account for this difference, we repeated the analysis with the baseline category logit model using only optimal-to-non-optimal variants as the COC category. Only DNA repair genes showed a significant shift towards being more intolerant (Mann-Whitney test p-value = 0.016), while the other two gene sets had no significant shift towards intolerance (Supplementary Figure 9). The AUCs also reflected the same overall pattern (Supplementary figure 10).

**Figure 3.**
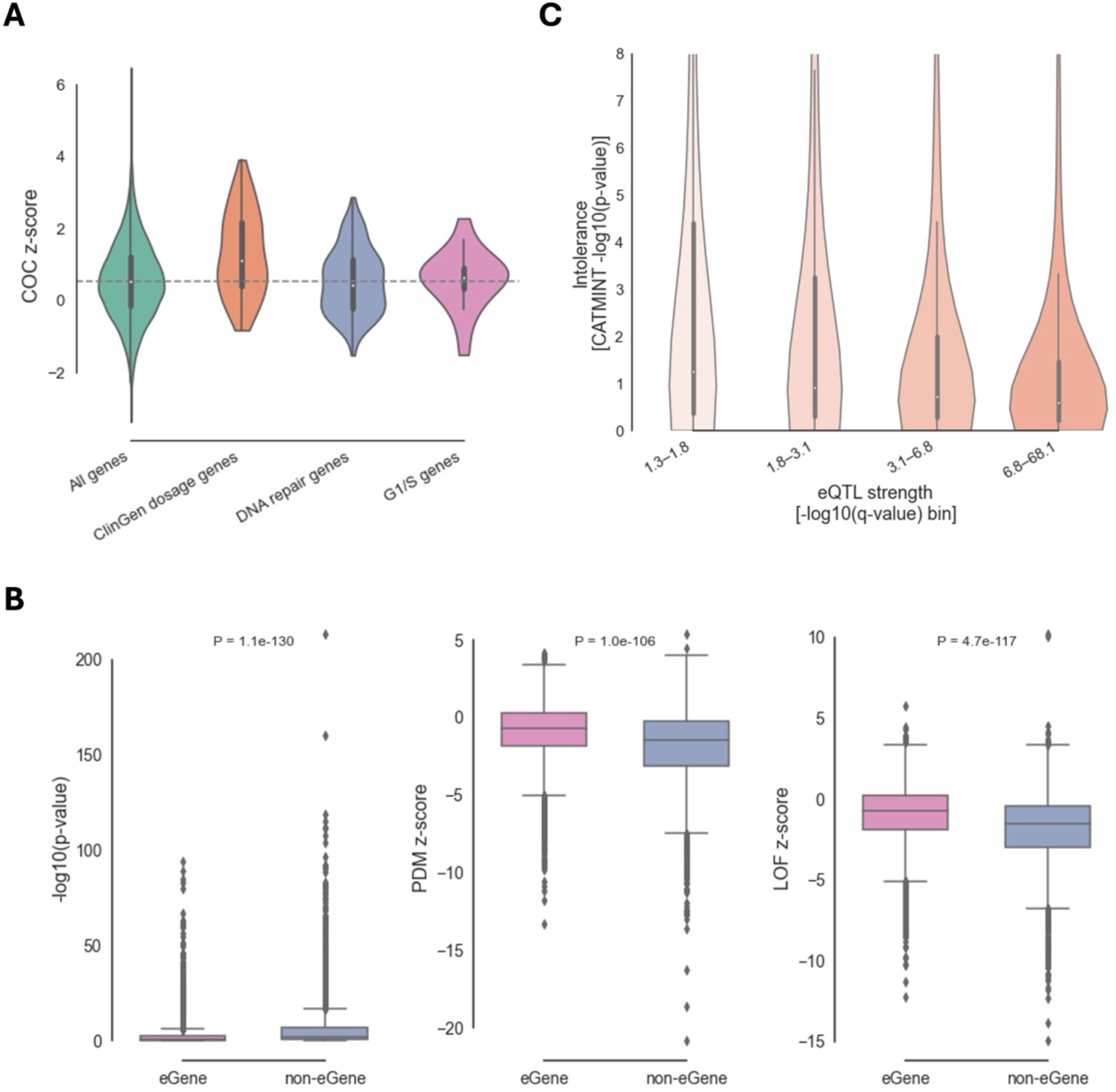
Assessing Intolerance Across Codon Optimality and eQTL-Linked Genes. (A) Distribution of intolerance z-scores for codon optimality–changing (COC) variants across all genes, ClinGen dosage-sensitive genes, DNA repair genes, and G1/S-phase genes. None of the sets showed a significant shift toward stronger constraint compared to all genes (one-sided Mann–Whitney U test p-values: ClinGen dosage = 1.0, DNA repair = 0.449, G1/S = 0.664), suggesting limited synonymous-level intolerance in these categories. (B) CATMINT intolerance scores comparison between eGenes and non-eGenes in brain frontal cortex. eGenes show significantly reduced intolerance across overall intolerance level (one-sided Mann–Whitney p = 1.1 × 10⁻¹³⁰), LOF z-scores (p = 4.7 × 10⁻¹¹⁷), and PDM z-scores (p = 1.0 × 10⁻¹⁰⁶). eGenes are more tolerant to genetic perturbation. (C) Intolerance (CATMINT –log₁₀(p-values)) distribution stratified by bins of eQTL significance (–log₁₀(q-value)) in brain frontal cortex among eGenes. eGenes with stronger regulatory signals (lower q-values) tend to be less intolerant.

**Figure 4.**
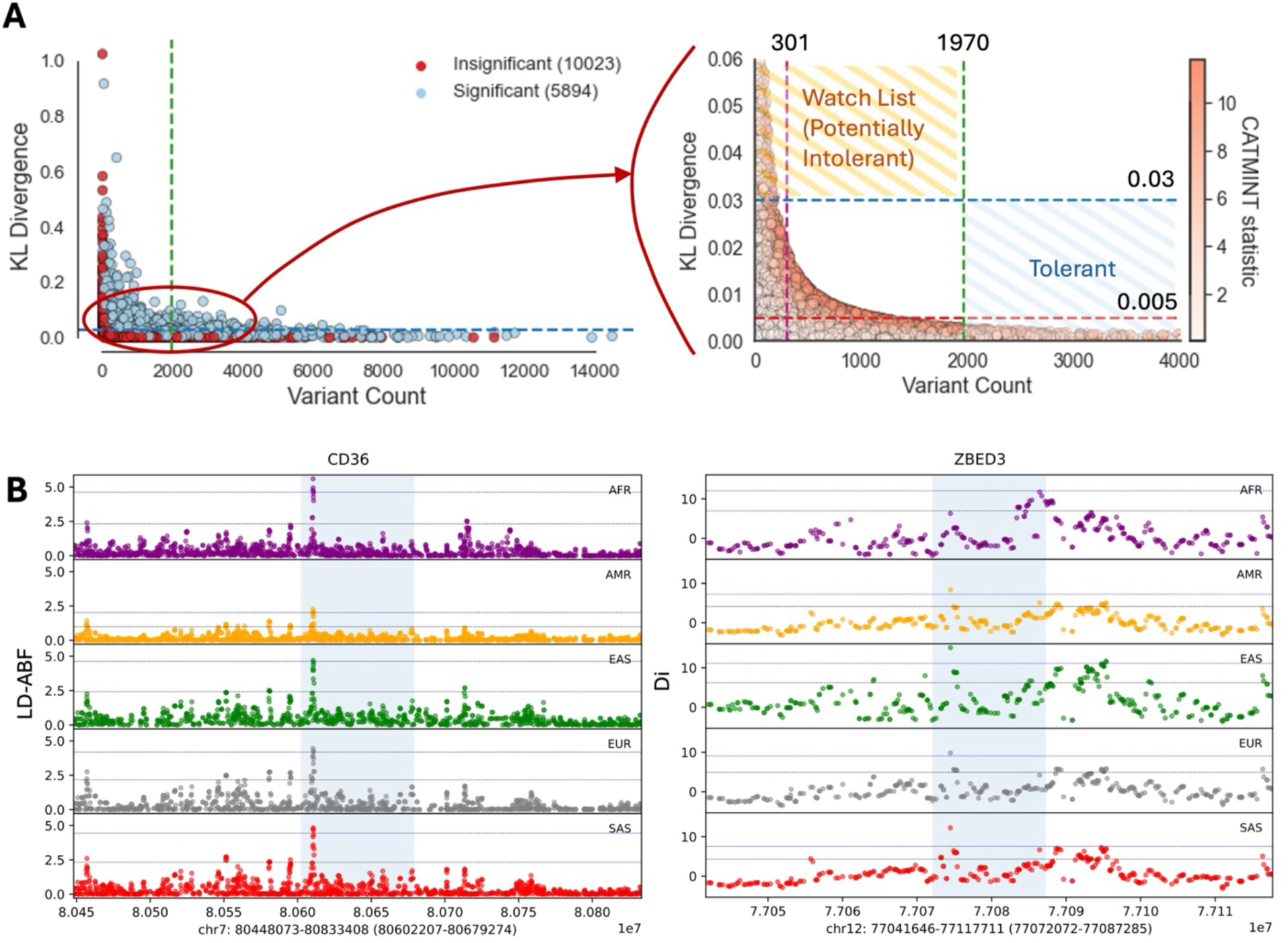
Identifying overlooked or selectively variable genes through power-aware modeling and population genetic scans. **(A)** KL divergence versus variant count for all genes analyzed by CATMINT. Genes with high KL divergence (> 0.03) but insignificant (they also have low variant counts) are classified as “potentially intolerant” or on the watch list, as their deviation from neutrality may be biologically meaningful but not statistically significant due to limited power. Genes with a large number of observed variants (>1970) yet insignificant in CATMINT (they also have small KL divergence values) are classified as “tolerant” genes. The zoomed-in panel focuses on non-significant genes, highlighting the watch list (upper left) and tolerant (bottom right) regions. Dotted lines indicate key thresholds, and points are colored by CATMINT significance status. (B) Genome-wide selection scan shows the top-scoring gene for LD-ABF (left) over the CD36 region and for Di (right) over the ZBED3 genic region. Tests are run across the 5 continental superpopulations in 1KGP. Top 1% (blue dotted-line) and 5% (black dotted-line) percentile thresholds were calculated for each superpopulation. The light-blue shaded region indicates gene region.

We hypothesize the remaining discrepancy between our results and the previous results could be due to the differences in modeling approaches: our method accounts for gene-specific neutral expectations, while the prior method compares optimal to non-optimal variants against total synonymous counts without adjusting for variation in sequence context across genes.

### Less Intolerance Observed in eGenes Compared to Non-eGenes

eQTLs are widely used in fine-mapping analyses to help prioritize candidate genes/variants and uncover regulatory mechanisms underlying complex traits and diseases. However, eGenes (genes associated with at least one significant eQTL) tend to have greater expression variability across individuals, which indicates their tolerance to dosage changes, or even to functional variants in general. Consistent with this idea, eGenes’ tolerance to loss-of-function (LOF) variants has been reported in previous studies (Lek et al., 2016; Mostafavi et al., 2023; Võsa et al., 2021; Wang & Goldstein, 2020).

To further investigate this pattern, we obtained tissue specific eGenes (qval < 0.05), significant eQTL pairs, and gene expression level data from GTEx (Lonsdale et al., 2013). Non-eGenes for each tissue were defined as genes with high expression level (TPM>3) but not included in the eGene set (Method). These gene sets were intersected with our intolerance analyses results. We identified a significant decrease in the overall intolerance levels (p-values from CATMINT) in eGenes compared to non-eGenes, across all tissue types. For example, in brain frontal cortex tissue, the Mann-Whitney test p-value reached 1.1e-130 (Figure 3B; Supplementary Data 4). This decrease in intolerance in eGenes was further identified in PDM variants (p-value = 1.0e-106) and LOF variants (p-value = 4.7e-117), with similar trends observed in other tissues (Figure 3B; Supplementary Data 4). To understand the quantitative relationship between intolerance and eQTL strength. We calculated correlations between CATMINT’s -log10(p-values) and -log10(q-values) of eGenes across all tissues, where q-values of eGenes indicate the statistical significance of regulatory association of each eGene. In brain frontal cortex, we observed a weak but significant negative correlation (Pearson’s r = −0.11, p-value = 2.6e-12; Spearman’s rho = −0.17, p-value = 4.6e-27), indicating that genes with stronger regulatory associations tend to be less intolerant. Similar patterns were observed in other tissues (Supplementary Data 4). To illustrate this trend, we stratified eGenes based on their eQTL association strength (based on q-values) in brain frontal cortex. Genes with stronger regulatory signals (lower q-values) showed higher CATMINT p-values, indicating reduced intolerance (Figure 3C). This observation is consistent with the negative correlation described above.

### Quantifying the Uncertainty in Intolerance Estimation to Avoid Overlooking Small Genes with Essential Functions

As with most constraint or intolerance scores, including CATMINT, the ability to identify intolerant genes depends on both the deviation between observed and expected variant compositions and the total number of observed variants in a gene, which influences the statistical power of the model. As a result, genes with more observed variants (typically larger genes) are more likely to reach significance compared to those with fewer variants-typically smaller genes. Even a substantial depletion consistent with true purifying selection may go undetected if the total number of observed variants in a gene is too low to provide sufficient statistical power. This limitation is common to other statistical models used to estimate gene intolerance.

To address this limitation, we further categorized non-significant genes into three groups: (1) likely tolerant genes, which have sufficient variants but show little deviation; (2) potentially intolerant genes, which show large deviation but insufficient variant counts to reach significance; and (3) uncertain genes falling between the two groups.

Because the CATMINT statistic in our framework (likelihood ratio test based on multinomial distribution) is equivalent to the weighted Kullbank-Leibler (KL) divergence (Supplementary section 10, Supplementary figure 11), we used KL divergence as a power-independent measure of deviation. We computed the cumulative distribution function of KL divergence for each essential gene set (Method; Supplementary section 11; Supplementary figure 12) and calculated the enrichment of highly deviated genes within each essential gene set relative to all genes (Supplementary figure 13-14). Based on patterns of enrichment in essential gene sets such as stillbirth genes and haploinsufficient genes, we chose thresholds of 0.005 and 0.03 for KL divergence. A KL divergence < 0.005 represents limited deviation (observed in 6.06% of stillbirth and 17.4% of haploinsufficient genes, compared to 43.5% of all genes), while a value > 0.03 indicated substantial deviation, observed in more than 40% of the essential gene sets (42.4% for stillbirth genes, and 41.1% for haploinsufficient genes) compared with only 12.2% in all genes (Supplementary section 11). These thresholds were chosen to help interpret deviation independently of statistical power.

To understand how statistical power influences the detection of intolerance, we conducted simulations to estimate observed variants (or sample size) required for a gene with a KL divergence of 0.005 to achieve a power of 0.95. Specifically, we simulated multinomial random variables across a range of sample sizes (1,000-4,000 variants), using alternative class probabilities designed to yield a fixed KL divergence at 0.005. To assess how power varies with sample size while holding the divergence from the null constant, we performed 1,000 simulations at each sample size. We found that genes with more than 1970 observed variants consistently yielded a power above 0.95. Based on this, we categorized non-significant genes with more than 1970 variants (all of them have a KL divergence below 0.005) as potential “safe genes” (326 genes, Supplementary Data 5). For genes that were not statistically significant but exhibited a KL divergence greater than 0.03 (313 genes, they also have small variant counts), we classified them as potentially intolerant genes. For these genes, simulations were performed to determine the number of variants required for CATMINT to achieve a 0.95 detection power (Supplementary Data 6).

These simulation results showed how statistical power limitations can lead to under-detection of intolerance in genes with fewer variants, often shorter genes, even when they show substantial deviation from neutral expectations. To investigate whether such genes may be overlooked in existing essential gene sets, we first compared the gene length distribution of commonly used essential gene sets. We found that many of these sets, including ClinGen haploinsufficient, stillbirth genes, showed a shift towards longer genes (Supplementary Figure 16). We then tested whether essential gene sets were enriched or depleted among genes with low variant counts (<301 and >10) and large deviation (KL > 0.03), using Fisher’s exact test. Several gene sets showed depletion among small genes, such as OMIM Recessive genes (OR = 0.250, p-value = 0.00712) and neurodevelopmental genes (OR = 0.539, p-value = 0.332). In contrast, MGI essential genes showed a trend of enrichment (OR = 1.39, p-value = 0.144) (Supplementary Table 7; Supplementary Figure 18). Similar results (Supplementary Table 8; Supplementary Figure 19) were observed when using an alternate group defined by gene length (<519 bp, variant count >10; threshold justified in Method) which is expected given the strong correlation between gene length and variant count (R² = 0.95, p < 2e–16; Supplementary Figure 17). The observed enrichment or depletion patterns across gene sets may reflect the distinct methods these gene sets were generated. MGI essential genes were identified through homozygous knockout experiments in mice and are less influenced by gene size, whereas OMIM and ClinVar-based sets were curated from peer-reviewed literature and may be more susceptible to power-related biases, such gene length.

### Rare Occurrence of Genes with an Excess of Functional Variants in Humans

While CATMINT p-value provides a measure of deviation from neutrality, it does not capture directionality. To identify whether this deviation reflects excess or depletion of functional variants, we measured the directional deviation for PDM and LOF variant compositions for each gene (Method). Among the 15,819 genes (11,551 high confidence), only 14 (all high confidence) showed a significant excess of functional variants after multiple testing corrections (Supplementary Data 7). This result reinforces the utility of the CATMINT p-value directly as a reliable measure of intolerance. Moreover, it suggests that most of the genes tolerant to missense or LOF variants, i.e. genes that did not show depletion of those two types of variants, may be experiencing just relaxed selection, or under minimal selection pressure (Lahti et al., 2009; Wertheim et al., 2015), rather than undergoing positive or balancing selection. However, to further distinguish these, additional analyses are needed, such as analyzing the allele frequency spectrum, haplotype structures, and enrichment or increase of frequencies on specific variants.

Among the 14 genes with an excess of functional variants, DNMT3A and TET2 are known genes that show evidence of clonal hematopoiesis of indeterminate potential (CHIP). These could serve as positive controls in our analysis. Somatic mutations, particularly LOF mutations, in CHIP genes accumulate with age and can be misclassified as germline variants in blood-derived sequencing data. This would then lead to an increase in functional variants, and result in these genes showing less functionally constrained in population datasets like gnomAD (Jaiswal et al., 2014; Karczewski et al., 2020). In our analysis, those genes would show significantly more elevated functional variant counts than expected. All 14 genes identified as having excess functional variants also showed low intolerance in other established constraint measurements (Supplementary Data 7).

To follow up on genes with significant excess of PDM and/or LOF, two genome-wide scans were conducted to detect balancing and positive selection across the 1000 Genomes Project (1KGP) Phase 4 (Fairley et al., 2020) data. These scans run for each superpopulation in 1KGP: European (EUR), East Asian (EAS), American (AMR), South Asian (SAS), and African (AFR), using the LD-ABF (Hayeck et al., 2024) and D_i_ statistics (Akey et al., 2010). The LD-ABF statistic identifies balancing selection by testing for elevated density of polymorphisms and linkage disequilibrium in a region. While the D_i_ statistic detects regional shifts in F_st_ between a target population and all others to pinpoint locus-specific divergence, such divergence at a given locus can be used to infer regions that have evolved due to local adaptation. Comparing signals across genes of interest to the overall genome revealed elevated LD-ABF in the CD36 and SMAD6 loci, and elevated D_i_ in the ZBED3, GJD3, CLIC6, TET2, and DNMT3A loci, all falling within the top 1% of scores for the respective statistics. These findings suggest increased mutability in these regions, which may correspond to genic regions with deviations in *z*_*PDM*_ and *z*_*L*0*F*_ scores but are not categorically under functional constraint.

Specifically, CD36 has been implicated in the pathogenesis of malaria due to its crucial role as one of the major receptors for p.falciparum-infected erythrocytes’ adherence to host endothelial cells. Variants that alter CD36 function or expression could affect this adherence process, potentially might influence an individual’s susceptibility to the disease (Aitman et al., 2000; Fry et al., 2009; Gao et al., 2023; Omi et al., 2003; Quintana-Murci, 2019), pointing to a potential selective advantage in regions where malaria is endemic.

## Methods

### Gene Model

We adopted the gene model from Ensembl (Martin et al., 2023)to ensure compatibility with the Variant Effect Predictor (VEP) tool we utilized for variant annotation (McLaren et al., 2016). We obtained the gene’s GTF file (Ensembl release-110) directly from the Ensembl’s ftp site (Data Availability). Our analysis focused exclusively on regions annotated with Consensus CDS (CCDS). In addition, pseudogenes were omitted from our analysis, resulting in a final count of 47,678 transcripts and 19,195 protein coding genes.

### Variant Annotation

We used Ensembl VEP (McLaren et al., 2016) to annotate single nucleotide variants (SNVs) as synonymous, missense, or loss-of-function (LOF). The LOF variants included splice acceptors, splice donor, start lost, stop gained and stop lost variants. Synonymous variants were further classified for their impact on codon optimality change (Pechmann & Frydman, 2013; Presnyak et al., 2015). The optimality score of each codon was adopted from a published study (Dhindsa et al., 2020), where the optimality of each codon was computed based on the abundance of the tRNA for each codon in human embryonic kidney 293T (HEK293T) cells. Positive scores indicate optimal codon (OP) while negative scores are non-optimal codons (NOP). Synonymous variants leading to a shift between OP to NOP were labeled codon optimality change (COC) variants. Synonymous variants that do not result in a codon optimality change were labeled codon optimality neutral (CON).

Missense variants underwent additional classification based on their predicted pathogenicity. Missense variants with polyphen2 (I. Adzhubei et al., 2013; I. A. Adzhubei et al., 2010) scores larger than 0.908 (defined by polyphen2) were labeled as probably damaging missense variants (PDM), otherwise they were labeled less damaging missense variants (LDM). To summarize, each possible variant was annotated as one of the five variant categories: loss-of-function (LOF), probably damaging missense (PDM), less damaging missense (LDM), codon optimality change (COC), or codon optimality neutral (CON).

### Extract gnomAD Variants within Coding Regions

The curated CDS region (generated in the first step) were intersected with the quality control (QC) files gnomAD exome version v4.1(Karczewski et al., 2020). We defined regions of high coverage in gnomAD v4.1 as those where at least 70% of the samples exhibit at least 10x coverage (Hayeck et al., 2019; Vitsios et al., 2021) based on the coverage file of gnomAD v4.1 exome. All exons required a minimum of 95% coverage with those gnomAD v4.1 high coverage regions. Regions with less than 5% overlap with the low complexity regions (LCR; Data Availability) or segmental duplicate regions (SegDup; Data Availability) were considered high confidence exon regions and were used in further analysis.

We further categorized transcripts into high-, medium-, and low-confidence groups based on exon-level annotation. Transcripts composed entirely of high-confidence exons were labeled as high confidence transcripts. To select a representative transcript for each gene, we prioritized the canonical transcript if it met high-confidence criteria. If not, we used the following strategy: The longest transcript that is high-confidence for each gene was selected; Genes not meeting the criteria for the high confidence group but containing at least one high confidence exon were placed in a medium confidence category; Transcripts with no high-confidence exons were initially excluded, but if they contained more than ten observed variants, they were revisited and included as low confidence transcripts.

The gnomAD exome v4.1 VCF files were intersected with high-confidence regions and low-confidence regions respectively. Variants in the high-confidence regions needed to pass the gnomAD internal quality criteria and be classified as PASS in the filter column as well as not labeled as segmental duplicates (SegDup). Variants in the low confidence regions, however, were only required to be classified as PASS in the filter column. Each variant was annotated according to the annotation file generated in the earlier step, and the number of observed variants in each one of the five categories were counted per transcript.

### Neutral Regions Extraction

To calculate the neutral variant rate for estimating the expected number of variants under neutrality, we first needed to isolate regions that were less likely to be directly under selective pressures. To approximate neutral regions, we excluded any sequences with known functional significance including gene regions, regulatory regions, and ultra-conserved elements. Also, we made sure to include only high-quality regions in the genome for the accuracy of our estimation by including only high coverage regions (at least 70% of the samples exhibit at least 10x coverage) of most updated whole genome date from gnomAD (gnomAD v3) and excluding segmental duplicates, repeats, and gaps (Supplementary Section 1; Supplementary Table 1).

### Neutral Variant Rates Calculation

The calculation of the neutral variant rate was based on the variants identified in the gnomAD whole genome v3.1 (Karczewski et al., 2020).Only high-quality variants passing gnomAD internal criteria (labeled as PASS and not SegDup) were included in the analysis. Our analysis involved implementing a 7bp sliding window across these high coverage neutral regions with a step of 1bp at a time. For each window, the variant at the central position was recorded and labeled by its 7-mer reference context and substitution type, i.e., ‘7-mer+nucleotide the focal site mutated to’. For example, ATCCAAGA denotes a C-to-A mutation at the central C in ATCCAAG (Supplementary Figure 1A). We systematically scanned through the neutral regions and tallied counts of each labeled mutation type observed in gnomAD v3.1 dataset. A variant rate for each label was computed based on the formula in Supplementary Figure 1B. Multiallelic variants are common (∼25%) in gnomAD v3.1 and should not be overlooked. Here, we treated them as multiple biallelic variants under the assumption that mutations at the same site occurred independently. To evaluate if our estimated neutral variant rates reflect the variation in the genomic sequences under neutrality, we selected several neutral regions matched to coding sequences in GC content and length and compared the estimated variant counts using our variant rate to the observed number of variants in the same regions in a number of datasets (Supplementary Section 3; Supplementary Table 2-5; Supplementary Figure 2).

### Expected Variant Counts Calculation

For the high, medium, and low confidence regions defined earlier, we computed the relative variant rate for every potential mutation at each genomic site (three mutations per site) based on the heptamer variant rate model we previously described. All possible mutations were annotated and the expected numbers of variants in each category were computed by summing up the variant rates for the mutations that fell within each category (Supplementary Figure 3).

### CATMINT: Likelihood Ratio Test (LRT) Model for Estimating Constraint

A likelihood Ratio Test (LRT) based on the multinomial distribution was applied to test how the composition of the variants deviated from that under neutral expectation. For a given gene, let 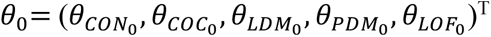 be the vector of multinomial cell probabilities under neutrality. Note that *θ*_0_ is a fixed parameter computed based on the estimated variant counts using the heptamer variant rate model we previously discussed. Let 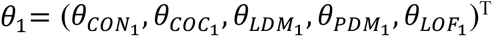 be a similar vector of free parameters that we are estimating. For the same gene, let 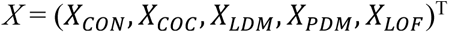 be the vector of observed variant counts in each category.

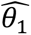 is the maximum likelihood estimate of *θ*_1_ given *X* under the multinomial model. A likelihood ratio is given by:

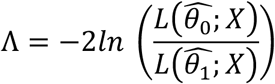

*L*(*θ*; *X*) is the multinomial likelihood function

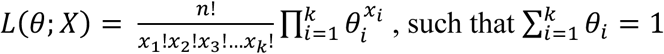

Where 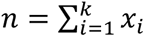 and k is the number of categories, in our analysis k = 5. Asymptotically, *T* will be distributed as 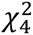 under the null hypothesis, i.e., *H*_0_: *θ*_1_ = *θ*_0_. The likelihood ratio test (LRT) was implemented in R (version 4.3.3) (R Core Team, 2024).

### Gene Sets for Overall Intolerance Validation

The OMIM (Online Mendelian Inheritance in Man) gene sets (Amberger et al., 2009) were extracted and refined to evaluate the predictive efficacy of the scores across multiple contexts. To extract these sets, we followed the approach detailed in the RVIS paper (Petrovski et al., 2013), using an updated version of the OMIM database (2023-03-03 update). We used the same search terms, i.e., “recessive”, “haploinsufficiency”, “dominant negative”, “de novo”, to query the OMIM database We additionally filter to genes with known sequence (denoted by “*” in OMIM) in known location (using “Gene Map Locus” in OMIM) and known phenotype (denoted by “+” and “Allelic variants” in OMIM). For the recessive genes, the list was manually curated so that the inheritance was AR only (remove AD, DR etc.). We also used RVIS’s (Petrovski et al., 2013) OMIM disease gene set and LIMBR’s (Hayeck et al., 2019) epilepsy and neurodevelopment genes directly.

In addition to the OMIM gene list, we used several publicly available gene sets curated by the MacArthur Lab (Data Availability), including Homozygous LoF tolerant genes (Lek et al., 2016), MGI essential genes (Blake et al., 2011; Georgi et al., 2013; Liu et al., 2013), ClinGen haploinsufficient genes (Rehm et al., 2015), and Olfactory receptors (Mainland et al., 2015). Both the Olfactory receptors genes and the homozygous LOF tolerant genes were used as genes that are known to be tolerant of variation. The stillbirth genes were expected to be the most constrained sets in our validation. Three gene sets were obtained from Stanley et al. 2020 including genes with molecular diagnosis previously associated with stillbirth (stillbirth_previous), genes previously associated with stillbirth but strong candidate with phenotype expansion (stillbirth_phenotype_expansion), and stillbirth candidate genes that were identified in a comparison of cases (stillbirth samples) and controls. The three stillbirth genes were grouped together to get the stillbirth_all gene set. A non-disease control set was derived by taking all genes that were not found in any of the gene sets described above.

### Variant Category Specific Intolerance

For each gene, we modeled the variant category probabilities, *θ*s, (defined in the LRT model above) using a baseline category logit model with codon optimality neutral variants designated as the baseline category. Mathematically, the model is expressed as:

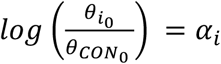

for the expected neutral variant probabilities, and

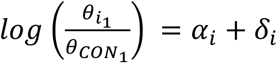

for the observed variant probabilities, where i represents the variant categories: COC, LDM, PDM, LOF. Pseudo-counts were added to all categories to address issues with zero counts i. For each gene, we used pseudo-counts across the five categories that are allocated in proportion to their expected composition. Thus, the pseudo counts function as a ‘prior’ for each gene, ensuring stability in the analysis.

The *δ* values capture changes in log-odds-ratio shift from neutrality for each variant category. The baseline category logit model was fitted using the ‘multinom ‘ function from the ‘nnet’ package in R (Venables & Ripley, 2002), which allowed for the specification of a multinomial family model with an offset. This offset is critical as it allows for adjustment based on the baseline log-odds provided by the neutral variant probabilities. From the model’s output, we extracted the *δ* values and their associated z-scores for the four categories - COC, LDM, PDM, and LOF.

### Enrichment of Variant Category Specific Constraint in Super Gene Families

We used the log-odds (*δ*) and z-scores estimated from the baseline category logit model described above to assess whether specific gene functional groups exhibit an enrichment of genic intolerance. Specifically, we analyzed the frequency of genes with elevated genetic intolerance (defined as those in the top 10% of z-scores) within each super gene families (SGFs), as defined by the Human Genome Organization (HUGO) Gene Nomenclature Committee (HGNC; Data Availability). For each SGF, the enrichment of constrained genes was then calculated for each category of variants with Fisher’s exact test. A higher enrichment score indicates greater intolerance to mutations within that category.

To identify SGFs with distinct profiles of intolerance, we computed relative enrichment ranks for each SGF across three variant categories: COC, PDM, and LOF. For each category, we assigned labels based on the ranking: High (H) - the rank was above 90th percentile, Low (L) - below the 10th percentile, and Medium (M) otherwise. Based on the H/M/L labels across the variant categories, we defined several enrichment patterns. SGFs that are labeled as H (for COC)-L (for PDM)-L (for LOF) were classified as COC-specific intolerant, while those labeled as L-H-H, or M-H-H were grouped as PDM and LOF intolerant. SGFs with patterns L-H-M or M-H-M were defined as PDM-specific intolerant, and those with patterns L-M-H, M-M-H as LOF-specific intolerant. Within each pattern group, we ranked SGFs by their corresponding enrichment value in the defining category (e.g., COC enrichment for COC-specific intolerant SGFs).

### Validation of Relative Intolerance between PDM and LOF Variants Using ClinVar Variant Patterns

ClinVar (Landrum et al., 2014) variants, with their corresponding genes, were downloaded (clinvar_20220910.vcf.gz; Data Availability). The variants were categorized into three groups based on their pathogenicity indicated by ClinVar: (1) Pathogenic: including both pathogenic and likely pathogenic SNVs; (2) Uncertain: including SNVs labeled with uncertain pathogenicity in ClinVar, and (3) Benign: including both benign and likely benign SNVs. For each category the number of missense variants and loss-of-function variants (nonsense; splice_acceptor_variant; splice_donor_variant; stop_lost) were counted for each gene (Pathogenic: *N*_*PM*_and *N*_*PL*0*F*_; Uncertain: *N*_*UM*_ and *N*_*UL*0*F*_; Benign: *N*_*BM*_ and *N*_*BL*0*F*_). The relative abundance of pathogenic missense variants over LOF variants was computed as *γ*_*P*_ = (*N*_*PM*_ − *N*_*PL*0*F*_)/(*N*_*PM*_ + *N*_*PL*0*F*_). Similar calculation was done for the benign (*γ*_*B*_ = (*N*_*BM*_ − *N*_*BL*0*F*_)/(*N*_*BM*_ + *N*_*BL*0*F*_)) and uncertain group (*γ*_*U*_ = (*N*_*UM*_ − *N*_*UL*0*F*_)/(*N*_*UM*_ + *N*_*UL*0*F*_)) as well.

To measure the relative constraint between the missense variants and loss-of-function variants, we computed the difference (*Z*_*PDM*−*L*0*F*_) between *Z*_*PDM*_ and *Z*_*L*0*F*_ (z-scores for probably damaging missense and loss of function, respectively) and scaled the by the variance of the difference. Specifically, we took 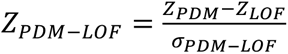 where 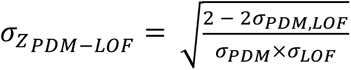 and *σ*_*PDM*,*L*0*F*_ is the covariance between the two. Genes with ClinVar variants were categorized into two groups based on the number of pathogenic missense and loss-of-function variants. Specifically, genes were assigned to one of the groups if the difference in the counts exceeded 10: *N*_*PM*_ - *N*_*PL*0*F*_ > 10 for ClinVar missense-leading group and *N*_*PL*0*F*_ − *N*_*PM*_ > 10 for ClinVar LOF-leading group. A Wilcoxon test was performed to compare the distribution of *Z*_*PDM*−*L*0*F*_ between the two groups of genes. A similar procedure was applied to the benign and uncertain ClinVar variants as our control tests, with some modification to the thresholds (Supplementary Section 7).

### Characterizing Intolerance Across Known CHD Genes and De Novo Variant-Enriched Genes from Clinical Trios

Testing was done with two different gene sets. The first gene set corresponds to congenital heart disease (CHD) genes with either loss-of-function (LOF) variants or missense variants as identified in a disease association whole exome sequencing (WES) study and a meta-analysis (Morton et al., 2022). These were further classified by their degree of evidence, i.e., definitive genes which are statistically significant, or candidate genes which display enrichment of variation in cases, but do not reach statistical significance.

The second gene set that was investigated included 533 proband-parent trios (mean coverage >= 60x) from the Birth Defects Biorepository (BDB) from the Children’s Hospital of Philadelphia. Raw variant calls in the VCF format were obtained from the BDB and de novo variants were identified using a set of heuristic filters. First, strict heterozygous variants were identified using the bcftools filter command to identify heterozygous variants in probands that passed the site quality filter (FILTER=PASS) and homozygous reference genotypes in the parents. Further filtering strategies included the depth filter (DP >=30), variant allele frequency filter (VAF between 0.2 and 0.8), and a cohort allele count (AC < 5). The de novo variants were then annotated using VEP and categorized into two groups of interest: those with at least one moderate-impact de novo variant in a CHD case, and those with at least one high-impact de novo variant in a CHD case. Definitions of moderate- and high-impact variants follow Ensembl’s variant effect prediction (VEP) criteria (Data availability)

We test for a shift in the distribution of each variant category z-score for genes in the gene set compared to a collection of baseline genes that are matched on gene length. More specifically, the baseline genes were assembled by matching each gene in the candidate set by selecting genes from the out-of-group gene pool that are within 5% of its coding domain sequence length. This matching process is repeated 1,000 times to compute an average z-score for the permuted baseline set. Finally, candidate gene z-scores are compared to the average z-scores of their permuted, size matched, baseline gene sets using a paired t-test.

### Testing Intolerance to Codon Optimality Changes in Selected Gene Sets

We took the three gene sets in Dhindsa et al., 2020 that were suggested to be potentially intolerant to codon optimality changes (COC): G1/S genes, DNA repair genes, and ClinGen dosage genes. We compared the distribution of the COC z-scores in each gene set to that of all genes using Mann-Whitney U test. Additionally, we calculated the area under the receiver operating characteristic curve (AUROC) to evaluate how well the COC z-scores separate the gene set from the background. To better match the definition used in synRVIS, we also repeated the analysis using a modified COC category that included only optimal to non-optimal substitutions (OP -> NO). In this analysis, the reference (CON) group was redefined to include synonymous variants except those classified as OP -> NO.

### Assessing Gene Intolerance Among eGenes and Non-eGenes Across Tissues

We downloaded the cis eQTL (v8) data from the GTEx portal, including eGene calls and all significant variant-gene associations per tissue (Data Availability). In addition, we retrieved gene expression level data, represented as median gene-level TPM per tissue (Data Availability) For each tissue, eGenes were defined using a q-value threshold of 0.05, as suggested by the GTEx portal documentation. Non-eGenes were defined as genes with TPM > 3 and q-value ≥ 0.05.

For each tissue type, we compared the distribution of CATMINT -log10(p-value), *z*_*PDM*_and *z*_*L*0*F*_ between eGenes and non-eGenes using a one-sided Mann-Whitney test. For each eGene, we extracted the smallest q-value of its associated eQTLs. After that, we performed both Pearson and Spearman tests to evaluate the correlation between the gene intolerance (CATMINT -log10(p-value)) and eQTL strength (GTEx -log10(q-value)). Finally, eGenes per tissue were binned into four groups based on their -log10(q-value). The distribution of their gene intolerance, *z*_*PDM*_ and *z*_*L*0*F*_ for each group were visualized using box plots.

### Assessment of Statistical Power for CATMINT

The power of CATMINT is influenced by both the sample size (i.e., the number of observed variant count) and the degree of deviation between the observed and expected compositions. We used Kullback–Leibler (KL) divergence to quantify this deviation. The CATMINT test statistic, based on the log-likelihood ratio, can be expressed as a function of the variant count n and KL divergence *D*_*KL*_ (see Supplementary Section 10 for detailed derivation; Supplementary Figure 11):

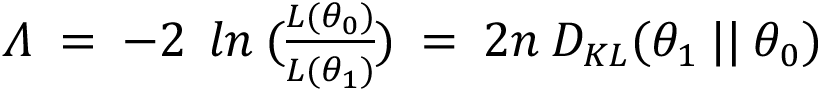

We calculated the KL divergence for each gene and generated empirical cumulative distribution functions (CDFs) for several representative gene sets described in Method (Supplementary Figure 12). To assess whether essential gene sets such as stillbirth or haploinsufficient genes show greater genetic deviation and to choose reasonable thresholds of large and small deviation, we compared their KL divergence distributions to that of all genes by subtracting the CDF of all genes from that of each gene set (Supplementary Figures 13 and 14). Additionally, we simulated 100 random gene sets of the same number of genes as the essential gene sets to confirm that the observed enrichment of genetic variation was not due to randomness (Supplementary Figure 15). KL divergence thresholds of 0.005 (small) and 0.03 (large) were chosen based on the CDFs, observed enrichments, and simulation results (Supplementary Section 10).

To estimate the number of variants needed to achieve a significance level of 0.05 for a given KL divergence (k), we used the observed and expected variant compositions of a gene with a KL divergence closest to k. We performed 1,000 simulations for each variant count within a reasonable range. For genes with KL divergence greater than 0.03 that were not identified as significant by the CATMINT (potential Intolerant genes), we calculated the statistical power, and the number of variants required to achieve a significance level of 0.03.

To assess whether essential gene sets are enriched in small, but substantially deviated genes (KL divergence > 0.03, and number of variants < 301 and > 10 for meaningful deviation), we counted the number of genes from each essential gene set that were in this small but substantially deviated gene group and those that were not, and we conducted Fisher’s exact tests. Since variant count is highly correlated with gene length (R² = 0.95, p < 2e-16; Supplementary Figure 17), we also defined small genes using gene length. We applied linear regression to gene length and observed variant count. Based on the regression, a gene length of 519 bp corresponds to 301 observed variants. We repeated the enrichment analysis for genes with length < 519 bp, KL divergence > 0.03, and variant count > 10 to ensure substantial and meaningful deviation.

### Identify Genes That Have More Functional Variation Than Their Neutral Expectation

To identify genes with an excess of segregating functional variation, we applied the categorical logit regression model described earlier and looked for evidence of an increase in the ratio of observed PDM or LOF variants over neutral expectation. To do so we calculated one-sided p-values for the positive tail of the z-scores for PDM, LOF or the sum of the two. The z-score of the sum of PDM and LOF constraint was computed using the formula below:

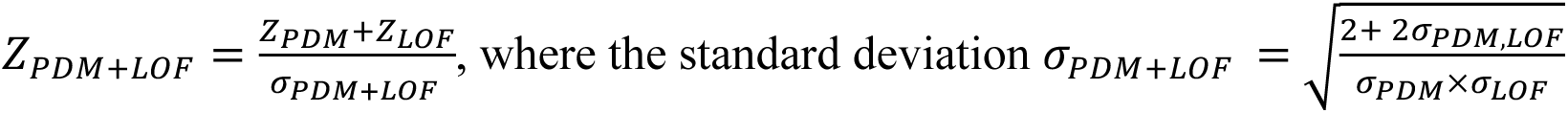

We then used the Cauchy combination test to integrate the three p-values into a single combined value. We then applied a Bonferroni correction to account for multiple testing.

### Genome-Wide Scan for Balancing and Positive Selection and Percentile Analysis

We analyzed whole-genome sequencing data from 2,003 unrelated individuals across five super-populations in the 1000 Genomes Project, excluding parents and filtering out low-complexity regions, blacklisted regions, centromeres, and telomeres (Supplementary Section 13).

To assess balancing and positive selection, we used two complementary statistics: LD-ABF and Di. LD-ABF, sensitive to more recent selection, tests for elevated levels of SNP density and linkage disequilibrium (LD) by fitting Bayesian logistic regression models between a target SNP and surrounding variants using logF priors. Di captures high-frequency divergence by quantifying locus-specific allele frequency differences between populations based on unbiased pairwise FST estimates.

Both LD-ABF and Di statistics were computed in 1,000 bp sliding windows. Variants were filtered using an Excess Heterozygosity Test, requiring a pExcHet > 0.0001 to be retained. For LD-ABF, a minor allele frequency (MAF) ≥ 0.05 in at least one population was required. For Di, a minimum MAF of 0.01 was applied across all variants. These thresholds ensured that only informative, high-quality variants contributed to the detection of selection signals.

Gene coordinates were obtained from NCBI using assembly GRCh38.p14 (GCF_000001405.40). For each population, the 95th and 99th percentile thresholds of LD-ABF and Di values were computed genome-wide and used to identify loci with elevated selection signals. We then checked whether the maximum statistic value within each gene of interest exceeded these thresholds and recorded their genome-wide percentiles.

## Discussion

In this study, we present CATMINT, a statistical model for estimating gene-level intolerance by jointly analyzing variant composition while accounting for sequence-context-specific mutability. Unlike previous models that rely exclusively on rare variants or de novo mutation rates, CATMINT incorporates both rare and common variants to estimate expected variant counts, enabling more comprehensive use of variant data. By modeling the shift in probability of variant class conditional on total variants, i.e. the variant composition, CATMINT reduces the impact of demographic biases that typically affect absolute mutation rates and variant counts, while remaining sensitive to relative depletion across functional categories. This compositional framework also has the advantage of implicitly accounting for local genomic features, such as recombination rate, methylation, and gene expression, that are typically modeled explicitly in mutation rate estimation.

A major advantage of our model lies in capturing variant category-specific intolerance patterns that reveal functional properties and mechanistic insights. This approach builds on the idea that natural selection shapes the variant tolerance of genes according to their functional roles. By identifying which types of variants are most strongly depleted in the population, we can infer which molecular features, such as protein structure, protein stability, dosage sensitivity, or regulatory precision, are most essential to a gene’s function. In this way, category-specific intolerance profiles offer an evolutionary perspective on the functional intolerance that characterizes each gene. To test this, we leveraged existing super gene family (SGF) definitions, which group genes based on sequence similarity and shared biological functions. Because SGFs reflect functional relationships, they provide a framework to assess whether category-specific intolerance patterns align with biological/functional properties. For example, potassium channel genes show strong intolerance to PDM variants while moderate to low intolerance to LOF variants, suggesting more sensitivity to structural alterations than to dosage changes. Calcium channel genes, however, are intolerant to both PDM and LOF variants. The difference between the two sets of channel genes could be explained by the higher redundancy of potassium channel genes where multiple genes can compensate for LOF in an individual gene (Chiang et al., 2017; Lazarenko et al., 2010), Calcium voltage-gated channel genes, by contrast, exhibit less redundancy and greater functional specificity, as they play critical roles in initiating precise and localized intracellular calcium signaling, which is essential for triggering tightly regulated cellular responses and the greater functional specificity of calcium channel genes for their critical roles for initiating precise and localized changes in the intracellular calcium ion level and thus triggering specific cellular responses (Hullin et al., 1993; Lauerer & Lerche, 2024; Liaqat et al., 2019).

Bromodomain containing (BRD) genes play key roles in reading epigenetic markers, regulating gene expression, and contributing to DNA repair and cell proliferation. Their intolerance to both LOF and PDM suggests that the expression and structure of these genes are precisely controlled, where disruptions can lead to significant cellular consequences. Notably, BRD proteins are often deregulated in cancer, with mutations in the bromodomains frequently observed across various tumor types (Bursch et al., 2024; Olley et al., 2018).

Olfactory receptor (OR) genes show intolerance to COC variants, despite their overall tolerance to nonsynonymous variants. OR genes are known to exhibit monoallelic and spatially restricted expression, requiring precise control of mRNA dynamics. Their intolerance to COC variants suggests the importance of regulated mRNA stability and translation efficiency to OR genes. Consistent with this, OR transcripts are shown enriched for AU-rich sequences, leading to reduced secondary structure, and more efficient translation. Also, OR genes are enriched with uORF, and unusually short 3′ UTRs, which affect post-transcriptional fate of the OR gene transcripts (Shum et al., 2015). More recent work suggests that many OR genes could produce longer 3′ UTR isoforms through alternative polyadenylation, indicating that their post-transcriptional regulation may be more complex (Doulazmi et al., 2019).

In addition to capturing the biological and functional properties of the genes, our model also has important clinical applications. In CES, variant interpretation typically relies on population frequency, predicted functional impact, and established gene-disease associations. This interpretation framework does not systematically account for gene-specific sensitivity to different types of mutations. Our results demonstrate that incorporating category-specific intolerance can provide additional biological context to improve diagnostic interpretation. Specifically, if a gene is highly intolerant to a particular variant type, observing such a variant in CES strengthens the likelihood it is pathogenic and related to the disease. Our analysis of ClinVar pathogenic variants supported this by showing that genes with stronger intolerance to PDM or LOF variants tended to carry more pathogenic variants of the corresponding type. Incorporating such gene-specific intolerance patterns into clinical workflows could improve variant interpretation and diagnostic decision making by prioritizing variants that align with a gene’s known sensitivity to mutation types.

Recently, ClinVar P/LP missense-to-LOF ratio has been used to prioritize genes with high therapeutic potentials, i.e., genes likely cause disease in gain of function mechanisms (Ressler & Goldstein, 2023). Although agreement between CATMINT’s relative intolerance and the ClinVar missense-to-LOF ratio is generally high, several discrepancies highlight strengths and limitations of the two approaches. Some genes with high ClinVar missense ratios do not appear more PDM-intolerant in our model. There are at least three possible explanations for this pattern. The first explanation is the bias in ClinVar and similar databases (such as OMIM) that they are enriched for well-studied genes and for variant types that are easier to interpret or more commonly tested. Such biases can distort gene-level P/LP missense-to-LOF ratios. In the ClinVar-based strategy, adding inheritance mode restriction and a low pLI (Lek et al., 2016) cutoff helps limit bias by restricting candidates to genes that appear LOF-tolerant from population genetics, providing independent support that a high missense ratio is not due to under-ascertainment of LOF variants.

A second explanation is that associated phenotypes may be late-onset or have little impact on fitness, so population data do not show strong selection against missense variants. Several late onset mendelian disease genes reflect this pattern. For example, missense mutations in MAPT cause frontotemporal dementia with parkinsonism (FTDP-17), a condition that typically emerges in mid- to late-adulthood, with an autosomal dominant inheritance (Hutton, 2001). ClinVar includes 23 P/LP missense variants, and only one P/LP pathogenic variant, while no strong intolerance/relative intolerance was identified for MAPT (*z*_*PDM*_ = −1.343, *z*_*L*0*F*_ = - 1.007, *z*_*PDM*−*L*0*F*_= −0.268). Similarly, pathogenic missense mutations in TTR (*z*_*PDM*_ = −2.186, *z*_*L*0*F*_ = −1.151, *z*_*PDM*−*L*0*F*_ = −0.830) destabilize the transthyretin tetramer and promote amyloid fibril formation, leading to hereditary transthyretin amyloidosis with adult-onset neuropathy or cardiomyopathy (Poli et al., 2023). Because these conditions present later in life, purifying selection in population datasets is limited, and relative intolerance appears weaker than the strong pathogenicity observed clinically (ClinVar P/LP missense: 85, ClinVar P/LP LOF: 0).

Finally, a third explanation is that pathogenic missense variants are clustered in specific domains, and thus PDM intolerance signal becomes diluted at the gene level. Examples include MACF1, KRT17, and KRT6A. MACF1 has ClinVar P/LP missense variants that cluster in the C-terminal GAR (Gas2-related) domain, a microtubule-binding region implicated in neurodevelopmental disease. In contrast, relative intolerance scores show much stronger LOF than PDM intolerance (*z*_*PDM*_ = −3.714; *z*_*L*0*F*_ = −14.946; *z*_*PDM*−*L*0*F*_= 8.793). For keratins, KRT17 and KRT6A also show weak gene-level intolerance, with slightly greater LOF intolerance (KRT17: *z*_*PDM*_ = 0.413; *z*_*L*0*F*_ = −0.433; *z*_*PDM*−*L*0*F*_= 0.695; KRT6A: *z*_*PDM*_= 0.642; *z*_*L*0*F*_ = −0.479; *z*_*PDM*−*L*0*F*_= 0.944), however, ClinVar P/LP missense variants are known to cluster in the terminal consensus domains (TEDs) of the α-helical rod. Specifically, KRT17 variants are mainly in TED01 (helix-initiation motif), while KRT6A variants occur in both TED01 and TED02 (helix-termination motif). Those domains are also among the most intolerant ones within each gene based on regional intolerance measurement (LIMBR; Hayeck et al., 2019). Such patterns highlight how domain-level missense clustering can be missed when intolerance is averaged across the entire gene and suggest that incorporating regional intolerance would improve prioritization.

We also see the reverse situation, where genes are PDM-intolerant, but ClinVar does not provide strong or consistent support. We also consider three explanations. The first is that some genes have too few P/LP variants in ClinVar to provide a reliable ratio for therapeutic prioritization. For example, POLD1 shows strong PDM intolerance (*z*_*PDM*_ = −5.839; *z*_*L*0*F*_ = - 0.282; *z*_*PDM*−*L*0*F*_ = −4.493), while ClinVar lists only five P/LP missense variants and no LOF variants. This shows how relative intolerance can reinforce ClinVar observations and help prioritize potential drug targets when clinical data are sparse. A second explanation is that the variant class recorded in ClinVar may not correspond to the true pathogenic mechanism. PPM1D shows stronger PDM than LOF intolerance (*z*_*PDM*_ = −4.326; *z*_*L*0*F*_ = 1.019; *z*_*PDM*−*L*0*F*_ = −4.641), while ClinVar reports nine LOF and no missense P/LP variants. The reported LOF variants are last-exon truncations that escape nonsense-mediated decay (NMD) and produce stable transcripts with gain-of-function effects. This mechanism is consistent with our relative intolerance results, even though the ClinVar annotation appears discordant, which shows that relative intolerance can sometimes capture the underlying biology more accurately than ClinVar counts. The third explanation is that pathogenic missense variants may be lethal very early in development and therefore absent from clinical databases. MIB1, a stillbirth-associated gene, shows much stronger PDM than LOF intolerance (*z*_*PDM*_ = −5.415; *z*_*L*0*F*_ = −5.734; *z*_*PDM*−*L*0*F*_ = −9.860), while ClinVar lists six LOF and only one missense P/LP variant. Mutations in MIB1 have been linked to left ventricular noncompaction (LVNC) and related cardiomyopathies, but the mechanism remains controversial. Early family studies reported one nonsense variant (e.g., R530X, LOF) and one missense variant with full penetrance for LVNC (Luxán et al., 2013), however, later mouse models found heterozygous R530X carriers were phenotypically normal (Siguero-Álvarez et al., 2023). These two variants are also present in gnomAD v4.1, though rare, potentially indicate tolerance. Dutch LVNC cohort analyses further identified three disease related truncating variants in MIB1, all co-occurring with TTN truncating variants, consistent with a modifying rather than primary role in causing LVNC (Mazzarotto et al., 2021). In such cases, relative intolerance metrics may provide additional information to help interpret discordant observations. These examples suggest that relative intolerance could provide complementary evidence to clinical databases: while ClinVar P/LP counts or ratios reflect accumulated clinical or research observations, relative intolerance provides a view from population genetics. Integrating these sources, and extending the relative intolerance metrics to regional level, will be an important future direction for clarifying controversial cases where the mechanisms remain uncertain.

We observed that eGenes tend to be less intolerant compared to non-eGenes, and this pattern was consistent across tissues. Possible explanations include: (1) greater expression variability among eGenes, making them more likely to be detected even with small sample sizes. And (2) the presence of their regulatory variants that are either more numerous or have larger effects, which would be detected as eQTLs. In contrast, genes where large expression shifts or regulatory variants are strongly selected against, are less likely to be detected as eGenes. Our findings, together with similar observations reported by several other groups (Lek et al., 2016; Mostafavi et al., 2023; Võsa et al., 2021; Wang & Goldstein, 2020) suggest that while eQTL-based fine-mapping approaches are valuable for linking regulatory variation to disease, they may preferentially capture genes that are more tolerant to expression variation and thus may not always overlap with the most dosage-sensitive or highly intolerant genes. Integrating intolerance information with eQTL signals may provide a more complete view of gene regulation and disease mechanisms.

Statistical models for estimating gene or region intolerance inherently rely on the ability to detect depletion of variation, which is directly influenced by gene size and the resulting statistical power. In most settings, genes that do not reach statistical significance are typically treated as less intolerant and excluded from further analysis. However, lack of statistical power to detect does not necessarily indicate lack of functional constraints. In our model, the power to detect intolerance depends on both observed variant counts and the deviation between observed and expected variant compositions. This introduces a potential bias: smaller genes, or genes under strong purifying selection with fewer observed variants, may fail to reach significance despite showing meaningful deviation. To address this, we explicitly classified non-significant genes based on their deviation magnitude (measured in KL divergence) and variant counts into three groups: (a) genes with large deviation but insufficient variants to achieve statistical significance (“watch list” genes), (b) genes with sufficient variants but minimal deviation from neutrality, and (c) genes falling between these extremes. Category (a) is particularly important, as these genes may represent biologically important but underpowered intolerant genes that are overlooked by standard analyses.

This speaks to broader biases that could also exist in the essential gene sets commonly used for validation of methods like the proposed one. Many curated gene sets, especially those drawn from clinical databases or literature-based sources like OMIM, may be biased toward longer genes that have more observed variants and are more likely to be identified in disease studies. These biases arise because longer genes have more opportunities for variants to occur, which increases their chances of being identified as essential or disease related genes in various gene prioritizing pipelines, such as de novo mutation studies, rare variant burden tests, GWAS. As a result, shorter genes, despite having potentially important functions, are underrepresented in these gene sets and often overlooked in downstream analyses. We evaluated enrichment within a subset of genes with low variant counts but high KL divergence in our model, indicating strong deviation from neutral expectation despite limited power (Supplementary Table 7-8; Supplementary Figure 18-19). Most established gene sets, including several OMIM-derived lists and sets related to neurodevelopment, were depleted in this group. In contrast, although MGI essential genes are not the most intolerant overall, they show the strongest enrichment among genes with large deviation but small variant counts. This difference potentially reflects that these sets are less influenced by gene length biases, likely due to their systematic experimental approaches (e.g., mouse knockout studies), while several other human disease gene sets are influenced by discovery and reporting biases favoring larger, better-studied genes. Together, these results highlight the importance of accounting for statistical power in interpretation when analyzing intolerance scores and suggest that future validation efforts may benefit from using more experimentally unbiased essential gene sets.

## Supporting information

Supplementary Doc

## Data Availability

Gene annotation: GTF gene annotation file was downloaded from Ensembl release 110 (https://ftp.ensembl.org/pub/release-110/gtf/homo_sapiens/Homo_sapiens.GRCh38.110.gtf.gz); Variant: gnomAD v2.1.1, v3.1.2, and v4.1.0 exome and genome variant datasets were downloaded from the gnomAD browser (https://gnomad.broadinstitute.org). 1kGP variants were downloaded from official FTP site (http://ftp.1000genomes.ebi.ac.uk/vol1/ftp/data_collections/1000G_2504_high_coverage/working/20220422_3202_phased_SNV_INDEL_SV/; release 2022-04). ClinVar variant annotations were obtained from NCBI ClinVar (https://www.ncbi.nlm.nih.gov/clinvar/; release 2022-09-10).

Region: LCR and SegDup regions were optained from resource indicated in gnomAD (https://gnomad.broadinstitute.org/help/vep). DNase I hypersensitive regions (open chromatin regions): Honeybadger2 release from the Reg2Map Project (https://personal.broadinstitute.org/meuleman/reg2map/HoneyBadger2_release/). Ultra-conserved elements (UCE) were downloaded from the UCE database (https://www.ultraconserved.org/). For the balancing selection analysis, LCR coordinates were obtained from https://github.com/lh3/varcmp/blob/master/scripts/LCR-hs38.bed.gz, ENCODE blacklist regions from https://github.com/Boyle-Lab/Blacklist/blob/master/lists/hg38-blacklist.v2.bed.gz, and centromere (acen) and telomere (gvar) positions from the UCSC Genome Browser cytoBand file (http://hgdownload.cse.ucsc.edu/goldenPath/hg38/database/cytoBand.txt.gz).

Gene set and gene family: Gene family annotations were retrieved from the HGNC database (https://www.genenames.org). Disease gene sets were obtained from OMIM (https://www.omim.org; accessed 2023-03-03) and manually curated based on published literature. Publicly available curated gene lists from the MacArthur Lab were downloaded from https://github.com/macarthur-lab/gene_lists.

Definition of moderate and high impact variants could be found in Ensembl (https://useast.ensembl.org/info/genome/variation/prediction/predicted_data.html).

GTEx data: cis-eQTL summary statistics and median gene-level expression data (TPM per tissue) from GTEx v8 were downloaded from the GTEx portal (https://gtexportal.org). All CATMINT gene-level scores generated in this study are provided in Supplementary Data 1, and all code used for statistical modeling and figure generation is available at (GitHub repositor; to be provided upon publication).

