## Supplementary Doc for "Functional Category-Specific Intolerance Reflects Genic Function and Clinical Relevance"

### 1. Defining Putatively Neutral Genomic Regions

To extract putatively neutral regions, we first excluded functional elements from the genome. Functional regions were defined by merging gene and regulatory annotations (include CTCF binding site/enhancer/open chromatin region/promoter/promoter flanking region/TF binding site) from Ensembl (Ensembl Biomart Version 103; hg38). DNase I hypersensitive regions were downloaded from Honeybadger2 in Reg2Map project (Data Availability). To exclude possible open chromatin regions, we chose the least stringent p-value threshold: 10e-2. These regions were originally in hg19 and converted to hg38 using UCSC liftOver (Hinrichs, 2006). Ultra-conserved elements (UCEs; Data Availability; McCormack et al., 2012) were downloaded with build hg19 and were similarly lifted over to hg38. Together, there were 2,028,545,510 bp of total functional regions after merging the genes, regulatory regions, DNase I regions, and ultra-conserved regions.

To mask potentially unreliable regions, we downloaded the repeat masked regions, segmental duplicates, and gaps from UCSC. Additionally, the relatively more recent repeats were extracted using repeatmasker (Smit et al., 2013) with -div 10 parameters. The total masked region (repeats and gaps combined) was 1,857,829,009 bp.

Finally, to ensure high confidence in variant detection, we restricted analysis to only high coverage regions in gnomAD v3.1, defined as the sites where more than 70% of the samples have at least 10x coverage. Intersecting with the criteria mentioned above, we had 294,308,993 bps of putatively neutral high-quality regions for calculating the neutral variant rate.

Supplementary Table 1 | Summary of Regions Used to Derive Putatively Neutral Sequences

| Region | Length |
|---|---|
| HighCov: High coverage regions (gnomAD v3) | 2,792,288,691 |
| RepeatMasker (recent repeats) | 305,608,415 |
| UCSC rmsk | 1,612,594,145 |
| UCSC Segdup | 910,687,638 |
| UCSC gap | 161,348,343 |
| All repeats + gap | 1,857,829,009 |
| Exclude all repeats/gaps from HighCov | 1,319,146,867 |

| | |
|---|---|
| Ultra conserved elements (UCE) | 240,391 |
| Ensembl gene + regulatory regions | 1,908,184,981 |
| HoneyBadger (10e-2) | 625,008,987 |
| All functional regions | 2,028,545,510 |
| HighCov neutral regions | 294,308,993 |

2. **Estimating Neutral Variant Rates**

To estimate context-dependent (heptamer) neutral variant rates, we scanned the high-coverage, putatively neutral regions (Supplementary Section 1) using a 7bp sliding window (step size = 1bp; Supplementary Figure 1A). For each window, variants observed at the central position were labeled by their 7-mer reference context and substitution type (check method for example). All possible 7-mers combined with 3 possible substitutions were considered, though not all possibilities (7-mer and substitution type) are present in the genome. The rate was calculated using the formula shown in Supplementary Figure 1B. Multiallelic sites were treated as multiple biallelic sites under the assumption of independent mutation events.

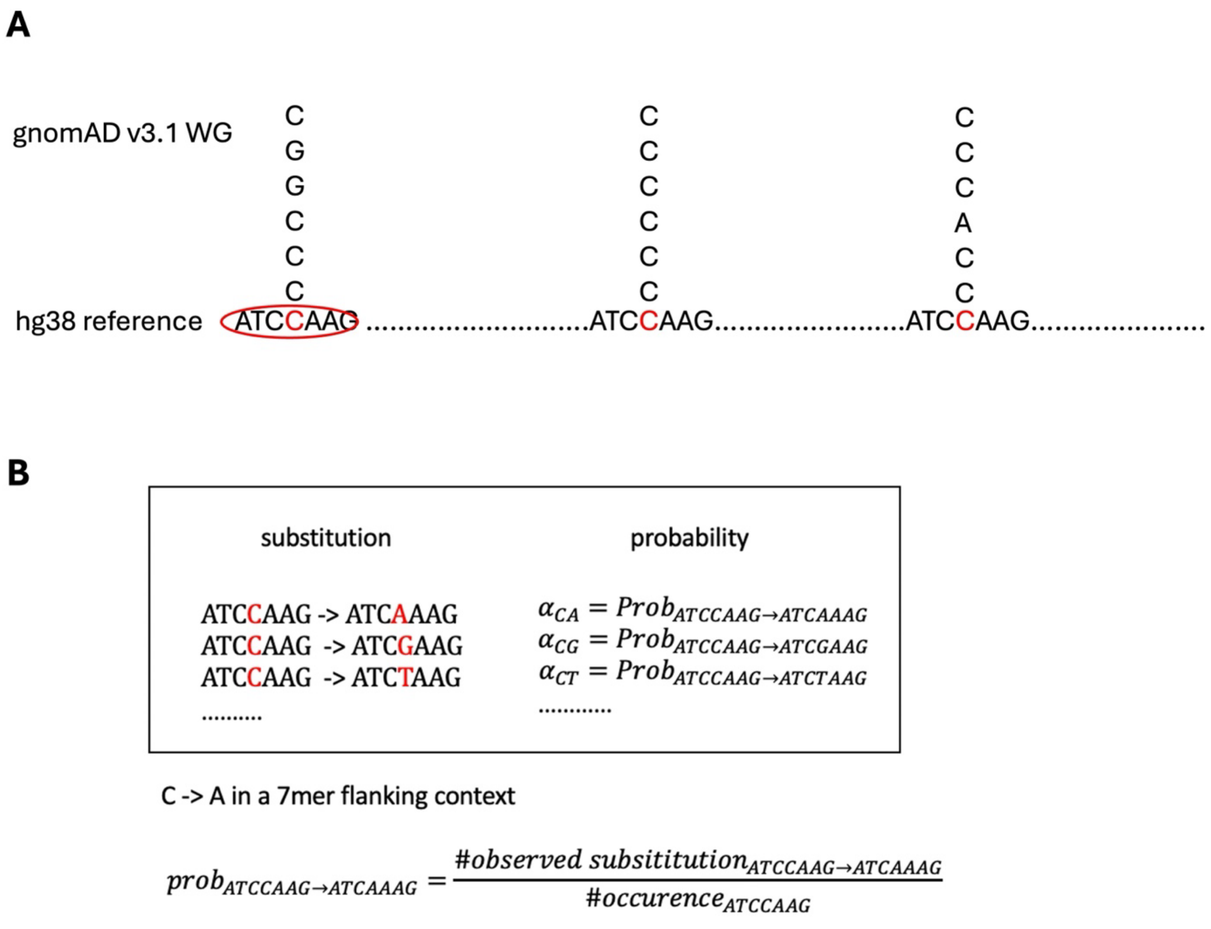


Supplementary Figure 1 | Estimating context-dependent neutral variant rate. (A) Schematic showing the 7bp sliding window used to identify variants and their sequence context in neutral regions. (B) Formula used to compute the variant rate per 7-mer substitution label, based on counts observed in the gnomAD v3.1 dataset.

### 3. Neutral Variant Rate Validation

We extracted sequences from coding regions (one transcript per gene, transcript selection detailed in Methods: Extract gnomAD Variants within Coding Regions) to calculate GC content and length for 17,725 transcripts. The matching process was performed using the R package - `nullranges` (Davis et al., 2023; Mu et al., 2023), which selected 17,725 neutral regions from a pool of 1,147,715 candidates. Regions that overlapped with the low-complexity and segmental duplicate regions (LCR and SegDup; Data availability; SegDup regions were lifted over to hg38) were also excluded from the candidates. The matching was based on the propensity score to match the GC content and length of the transcripts to ensure comparability.

For each selected neutral region, we calculated the cumulative variant rate across the entire region and used it as the expected variant count. Observed variant counts were obtained from the gnomAD v3.1 WGS dataset. We evaluated prediction accuracy by correlating observed counts with both transcript length and expected counts. Across all regions, both correlations are substantial, with expected counts based on variant rates showed stronger correlation with observed counts than did region length alone (Pearson's correlations: length - 0.957, mutation rate - 0.963). To further assess model performance, we further categorized the neutral regions into four length-based groups: 200bp-500bp, 500bp-1kb, 1kbp-2kb, and 2kb-3kb and repeated the correlation analysis. As shown in Supplementary Table 2, expected variant counts consistently outperformed length in all bins, indicating the improvement in the prediction accuracy when adding the variant rates.

To address the potential data leakage concerns from using the same dataset for estimation and validation (gnomAD dataset were used for neutral rate estimation and then for validation), we conducted follow-up analyses using the 1000 Genomes Project (1KGP; Data Availability). While 1KGP is independent of gnomAD v2.1, it was later incorporated into gnomAD v3.1. We therefore repeated the variant rate estimation using gnomAD v2.1 and performed validation using 1KGP. This approach ensured that the evaluation was conducted on a dataset entirely independent of the one used for rate estimation.

We extracted and counted number of variants per region from 1KGP and performed correlation tests between the observed counts and both region length and expected counts. For the gnomAD V3.1 mutation rate, high accuracy was observed across the full set of regions

(Pearson's correlation: length – 0.924; mutation rate – 0.940) as well as within subsets of regions defined by length ranges (Supplementary Table 3). The validation for gnomAD V2.1 mutation rate using 1KGP is shown in Supplementary Table 5 (see also Supplementary Table 4 for correlation results using gnomAD v2.1 observed variant counts; corresponding hg19 high coverage neutral regions were used). Supplementary Figure 2 further shows that expected variant counts estimated from gnomAD v2.1 and v3.1 are highly correlated (Pearson's correlation > 0.99 across both hg19 and hg38 neutral regions), supporting the robustness of our estimates.

Supplementary Table 2 | Correlation Results for Variant Counts for Different Length Ranges (gnomAD V3.1, w/ V3.1 rate)

| Length range | Num of regions | Cor w/ length | Cor w/ expected counts |
|---|---|---|---|
| (200,500) | 3064 | 0.58 | 0.71 |
| (500,1k) | 2618 | 0.42 | 0.55 |
| (1k,2k) | 4012 | 0.68 | 0.74 |
| (2k,3k) | 708 | 0.58 | 0.61 |

Supplementary Table 3 | Correlation Results for Variant Counts for Different Length Ranges (1kGP, w/ V3.1 rate)

| Length range | Num of regions | Cor w/ length | Cor w/ expected counts |
|---|---|---|---|
| (200,500) | 3064 | 0.47 | 0.61 |
| (500,1k) | 2618 | 0.44 | 0.62 |
| (1k,2k) | 4012 | 0.48 | 0.61 |
| (2k,3k) | 708 | 0.47 | 0.56 |

Supplementary Table 4 | Correlation Results for Variant Counts for Different Length Ranges (gnomAD V2.1, w/ V2.1 rate, using V2.1 neutral regions)

| Length range | Num of regions | Cor w/ length | Cor w/ expected counts |
|---|---|---|---|
| (200,500) | 2910 | 0.62 | 0.71 |
| (500,1k) | 2004 | 0.58 | 0.66 |

| | | | |
|---|---|---|---|
| (1k,2k) | 1846 | 0.66 | 0.74 |
| (2k,3k) | 1341 | 0.52 | 0.66 |

Supplementary Table 5 | Correlation Results for Variant Counts for Different Length Ranges (1kGP, w/ V2.1 rate)

| Length range | Num of regions | Cor w/ length | Cor w/ expected counts |
|---|---|---|---|
| (200,500) | 3064 | 0.47 | 0.62 |
| (500,1k) | 2618 | 0.44 | 0.64 |
| (1k,2k) | 4012 | 0.48 | 0.63 |
| (2k,3k) | 708 | 0.47 | 0.56 |

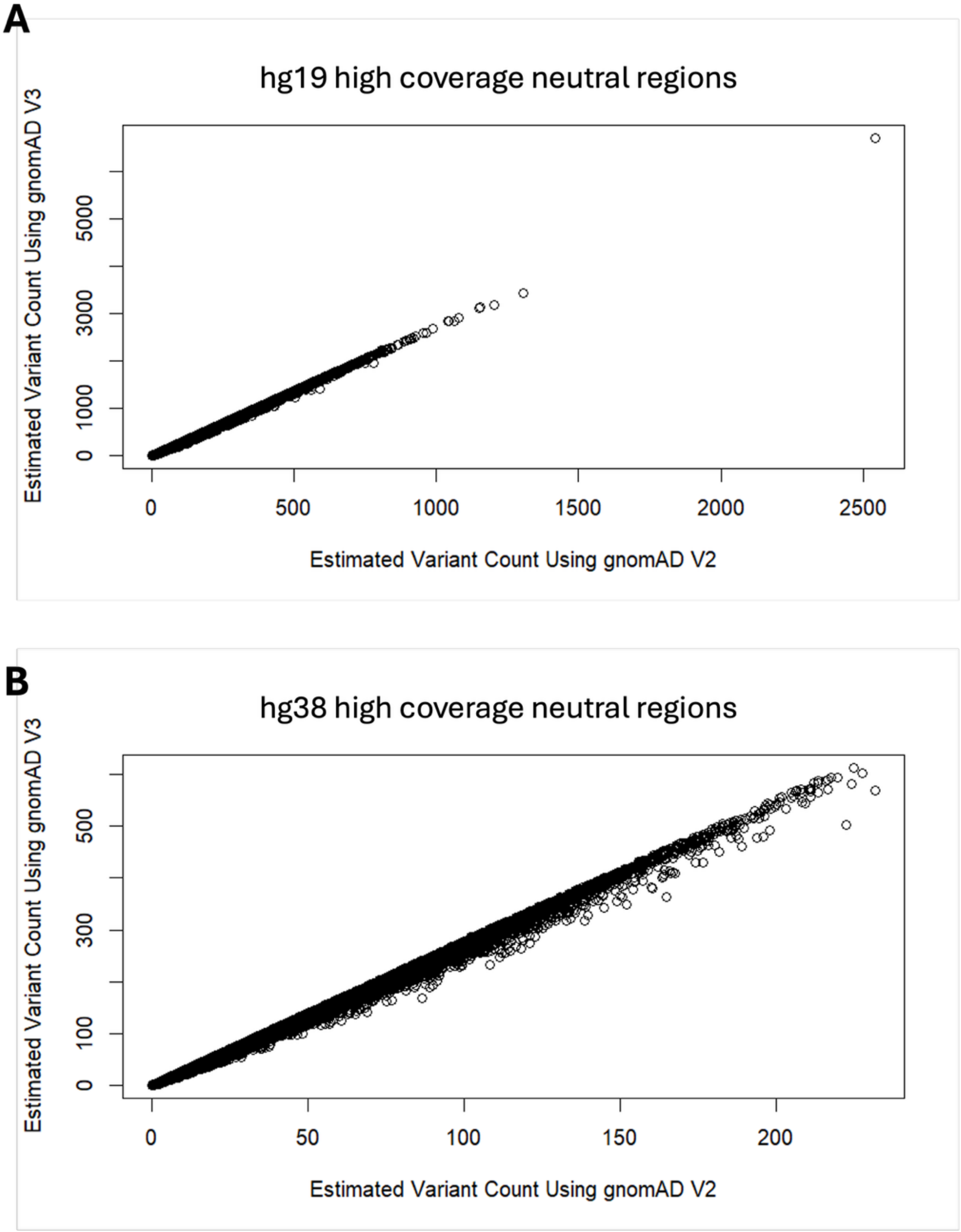


Supplementary Figure 2 | Consistency of Estimated Neutral Variant Counts Between gnomAD v2.1 and v3.1 variant rate. (A) Comparison of estimated neutral variant counts per hg19 high-coverage neutral region using gnomAD v2.1 (x-axis) and gnomAD v3.1 (y-axis) variant rates. Each point represents a matched neutral region. (B) same analysis performed on hg38 high-coverage neutral regions. In both (A) and (B), Pearson's correlation exceeds 0.99.

4. **Expected Variant Counts Calculation**

For each site, we considered all three possible substitutions and assigned a mutation rate using our 7-mer context-specific neutral model (see Supplementary Section 2). Mutations were then grouped into five functional categories (CON, COC, LDM, PDM, LOF) according to the gnomAD VEP annotations. For each transcript, the expected number of variants per category was calculated by summing the rates of mutations assigned to that category (Supplementary Figure 3). This allowed us to obtain transcript- and category-specific expectations that account for both sequence context and functional annotation.

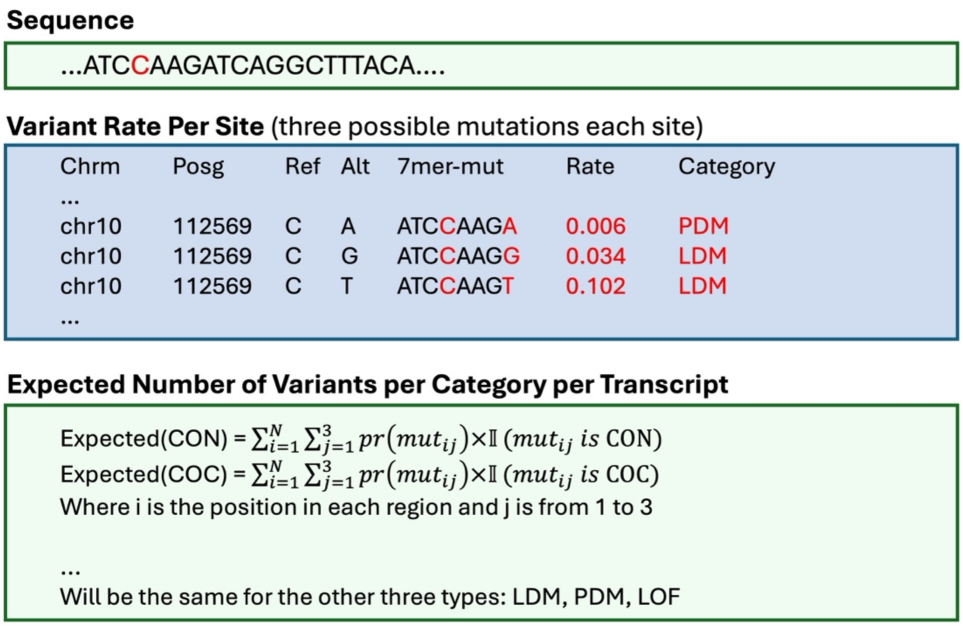


Supplementary Figure 3 | Illustration of context-dependent expected variant rate estimation per transcript. For each position in a sequence, the three possible substitutions are assigned rates based on their 7-mer context. These are then grouped by functional category (e.g., PDM, LDM), and the expected count for each category is computed by summing across sites.

### 5. Evaluation of the Gene Overall Functional Constraints

To evaluate the performance of CATMINT scores in in capturing gene level intolerance, we generated Receiver Operating Characteristic (ROC) curves and calculated the area under the curve (AUC) for each curated gene set. For this analysis, genes from essential, disease-related or tolerant sets were labeled as cases, while genes from a non-disease control set were labeled as controls (gene set selection could be found in method: Gene Sets for Overall Intolerance Validation). The ROC curves were plotted by comparing these case-control labels, with the prediction of intolerance informed by the CATMINT scores (i.e., p-values derived from likelihood ratio test (LRT)), with smaller p-values indicating stronger intolerance (Supplementary Figure 4).

To quantify enrichment, we ranked all genes by CATMINT scores and computed the proportion of genes from each gene set falling within the top intolerant percentiles: 1%, 5%, 10%, 15%, 20%, 25%, 30%, 40%, 45%, 50% based on the p-value ranks. Enrichment fold (EnF) was then defined as the observed proportion within a percentile threshold divided by the genome-wide background proportion (Supplementary Figure 5).

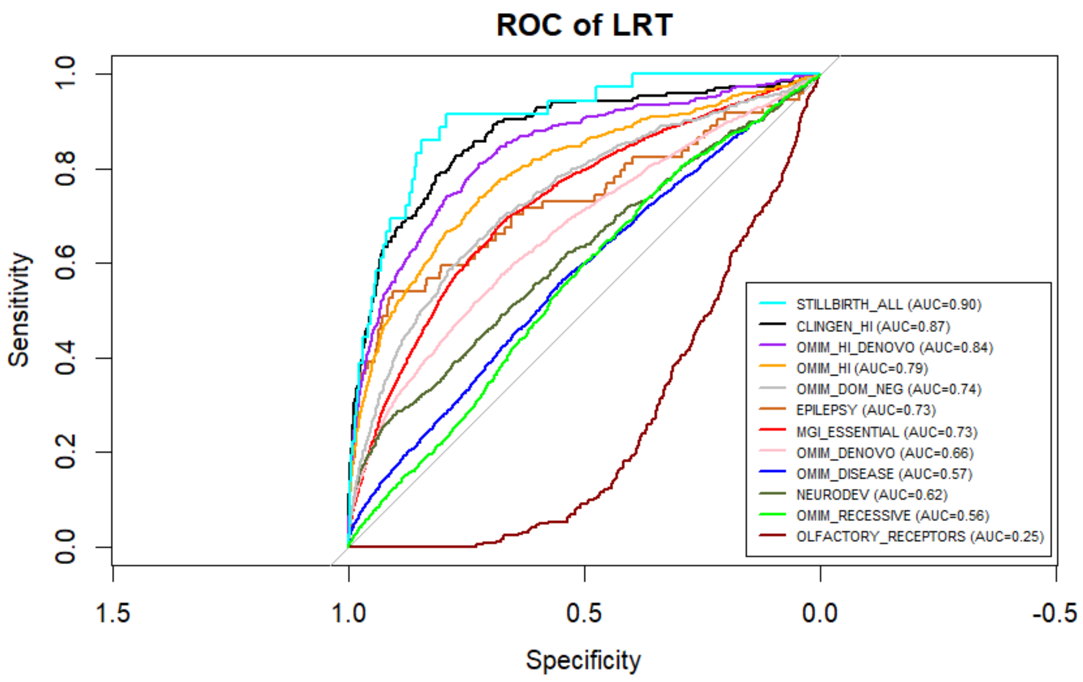

Supplementary Figure 4 | ROC of CATMINT using curated gene sets. ROC curves showing the performance of CATMINT scores in distinguishing essential and tolerant gene sets from control genes. Area under the curve (AUC) values are shown for each gene set.

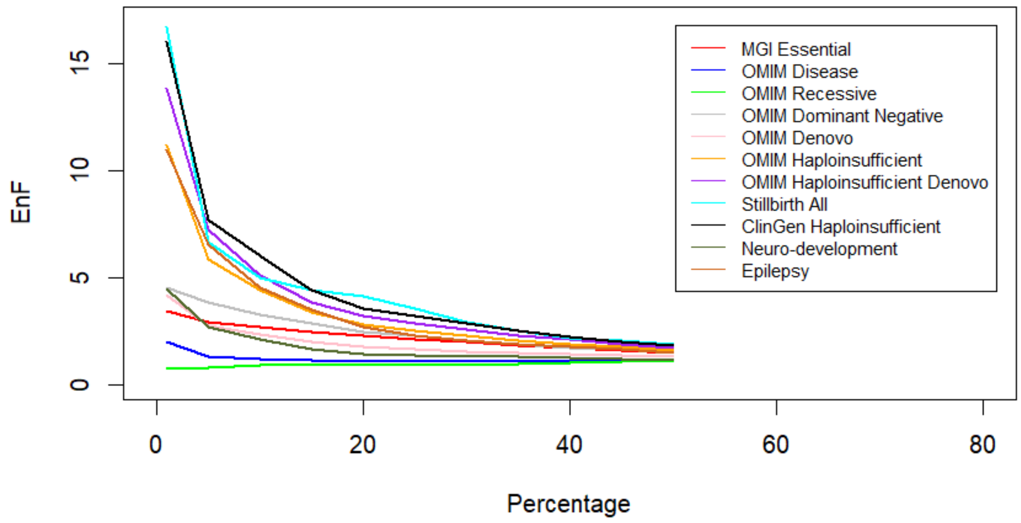


Supplementary Figure 5 | Enrichment of top intolerant genes in each essential gene set. Each line shows the enrichment fold (EnF) of a gene set at varying thresholds of top intolerant genes, defined by CATMINT scores. Enrichment is highest at stringent thresholds (e.g., top 1%), indicating that functionally important genes are strongly prioritized by CATMINT.

### 6. Compare CATMINT with Other Established Intolerance Scores

We compared CATMINT to several widely used gene-level intolerance scores, including Residual Variation Intolerance Score (RVIS), pLI, pNull, pRec, missense_z and LOEUF, and constraint_z (Chen et al., 2022; Karczewski et al., 2020; Petrovski et al., 2013). To ensure consistency and fair evaluation, we recalculated the RVIS using the gnomAD V4.1 whole exome data since the original RVIS, published in 2013, was based on a much smaller dataset. The quality control of the regions and variants was conducted identically to the procedures applied in our CATMINT analysis. Following the original RVIS framework, we performed a linear regression of the number of functional variants on the total number of variants per gene and used the studentized residuals as the constraint metric. The other four constraint scores: pLI, pNull, missense_z, loeuf were downloaded from the supplementary dataset 11 from (Karczewski et al., 2020). Descriptions of each metric can be found on the gnomAD constraint documentation page (https://gnomad.broadinstitute.org/help/constraint).

Supplementary Table 6 | AUC of Constraint Metrics on Validated Gene Sets

| Gene Set | RVIS | CATMINT | pLI | pNull | missense_Z | loeuf |
|---|---|---|---|---|---|---|
| Stillbirth (all) | 0.83 | 0.90 | 0.86 | 0.90 | 0.83 | 0.89 |
| ClinGen haploinsufficient | 0.77 | 0.87 | 0.85 | 0.87 | 0.76 | 0.87 |
| Stillbirth (previous) | 0.84 | 0.86 | 0.70 | 0.81 | 0.77 | 0.80 |
| OMIM haploinsufficient de novo | 0.78 | 0.84 | 0.78 | 0.83 | 0.76 | 0.83 |
| OMIM haploinsufficient | 0.73 | 0.79 | 0.74 | 0.79 | 0.72 | 0.79 |
| OMIM dominant negative | 0.71 | 0.74 | 0.68 | 0.74 | 0.73 | 0.75 |
| Epilepsy | 0.75 | 0.73 | 0.66 | 0.73 | 0.75 | 0.73 |
| MGI essential | 0.68 | 0.73 | 0.67 | 0.73 | 0.67 | 0.73 |
| OMIM de novo | 0.65 | 0.66 | 0.58 | 0.67 | 0.65 | 0.68 |
| Neuro developmental | 0.61 | 0.62 | 0.52 | 0.64 | 0.64 | 0.65 |
| OMIM disease | 0.57 | 0.57 | 0.47 | 0.61 | 0.60 | 0.62 |
| OMIM recessive | 0.54 | 0.56 | 0.40 | 0.57 | 0.55 | 0.58 |

7. **Evaluating PDM and LOF Intolerance Against ClinVar Pathogenic, Benign, and Uncertain Variant Groups**

To ensure that the observed shift in relative intolerance scores between genes with more ClinVar pathogenic missense versus LOF variants is not a false positive, we repeated the analysis using ClinVar benign and uncertain variants. Due to the limited number of genes with 10 or more LOF variants than missense variants in the benign and uncertain groups, we relaxed the criteria to include genes with more LOF than missense variants (Supplementary Figure 6B-C). To ensure a fair comparison, we also repeated the pathogenic analysis using this looser threshold (in addition to the original “10+ more missense” group) and included it here for reference (Supplementary Figure 6A).

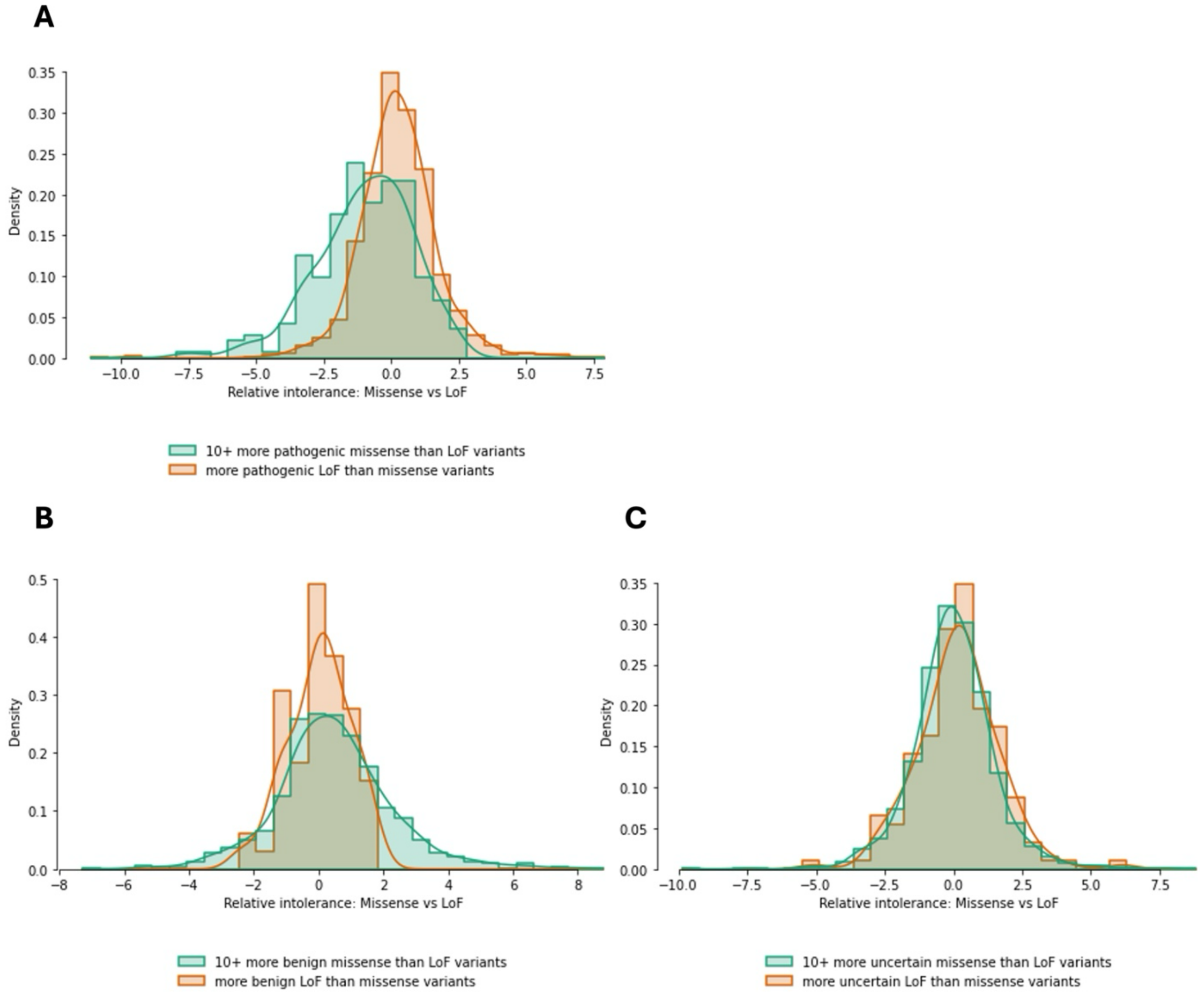

Supplementary Figure 6 | Control analysis to validate relationship between relative intolerance and ClinVar variant burden. (A) Pathogenic variants (10+ more missense vs. LOF, and more LOF vs. missense), (B) Benign variants (same criteria), and (C) Uncertain variants (same criteria). This analysis serves as a control to ensure that the signal observed in pathogenic variants is not an artifact of gene-specific biases in ClinVar annotations.

### 8. Using Relative Intolerance to Evaluate Gene's Therapeutic Potential

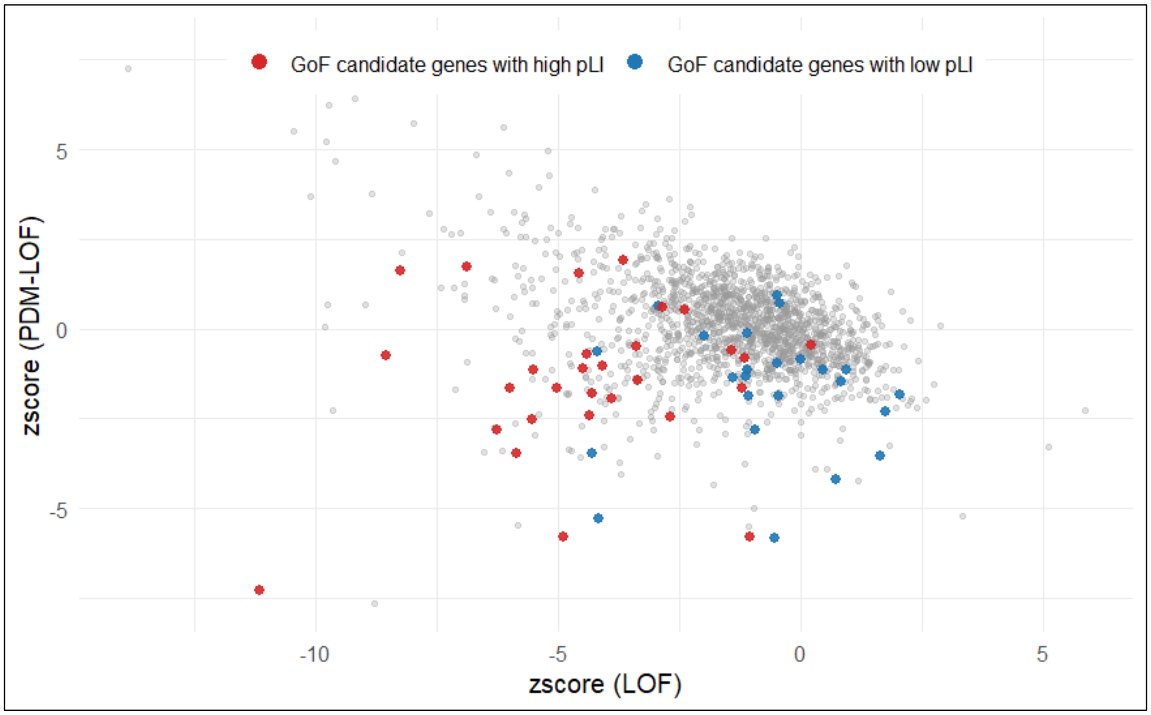


Supplementary Figure 7 | LOF intolerance versus relative intolerance (PDM – LOF) for several gene sets. Grey dots represent all OMIM genes. Blue dots represent potential gain-of-function genes, identified by a high ClinVar P/LP missense-to-LOF ratio and low pLI, suggesting they may be more suitable for therapeutic targeting. Red dots represent potential gain-of-function genes with high pLI.

### 9. Intolerance In Codon Optimality Change (COC) Variants

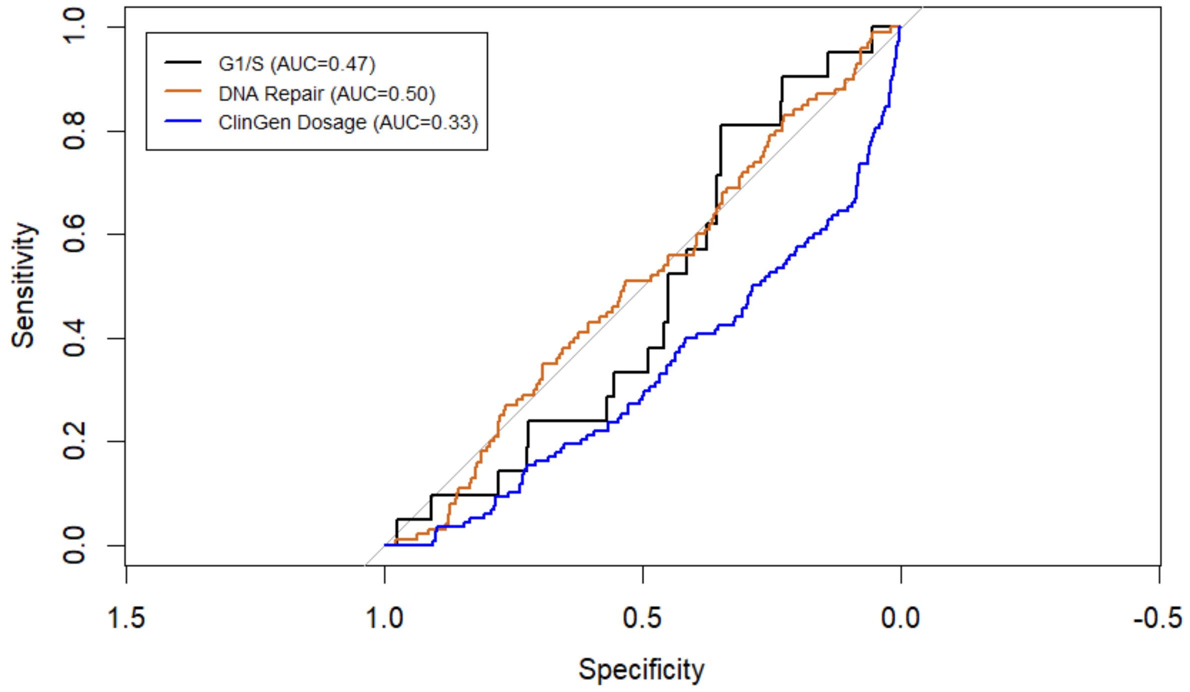


Supplementary Figure 8 | ROC curve with COC intolerance (z-score) for multiple gene sets. COC intolerance z-scores showed limited predictive power for G1/S, DNA repair, and ClinGen dosage-sensitive genes, with AUCs near or below 0.5.

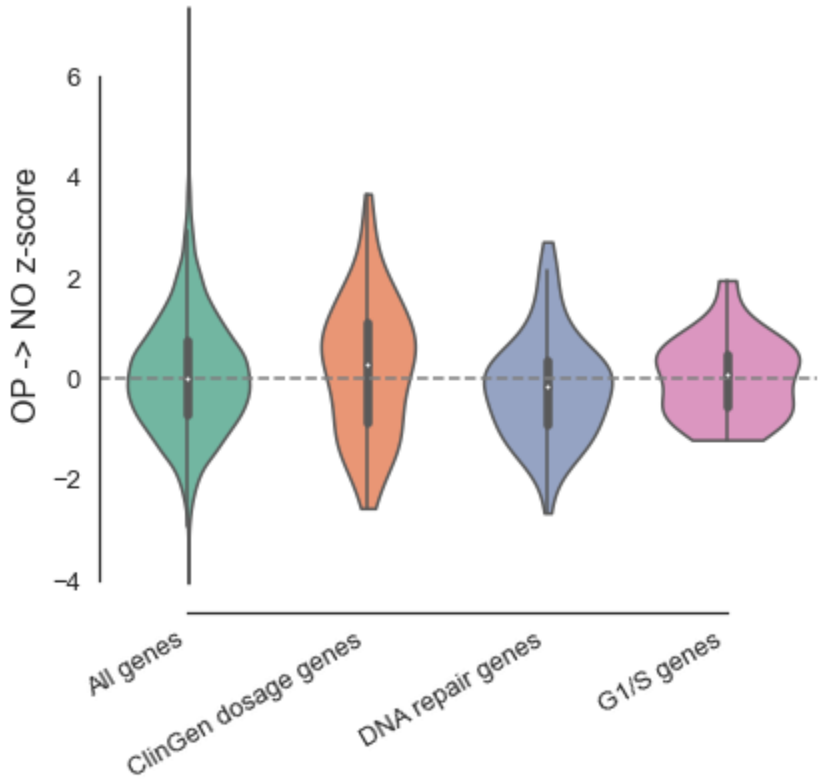


Supplementary Figure 9 | OP -> NO z-zcore distribution for multiple gene sets. Only DNA repair genes showed significantly higher intolerance than all genes (Mann–Whitney p = 0.016), while ClinGen (p = 0.945) and G1/S (p = 0.496) showed no or opposite shift.

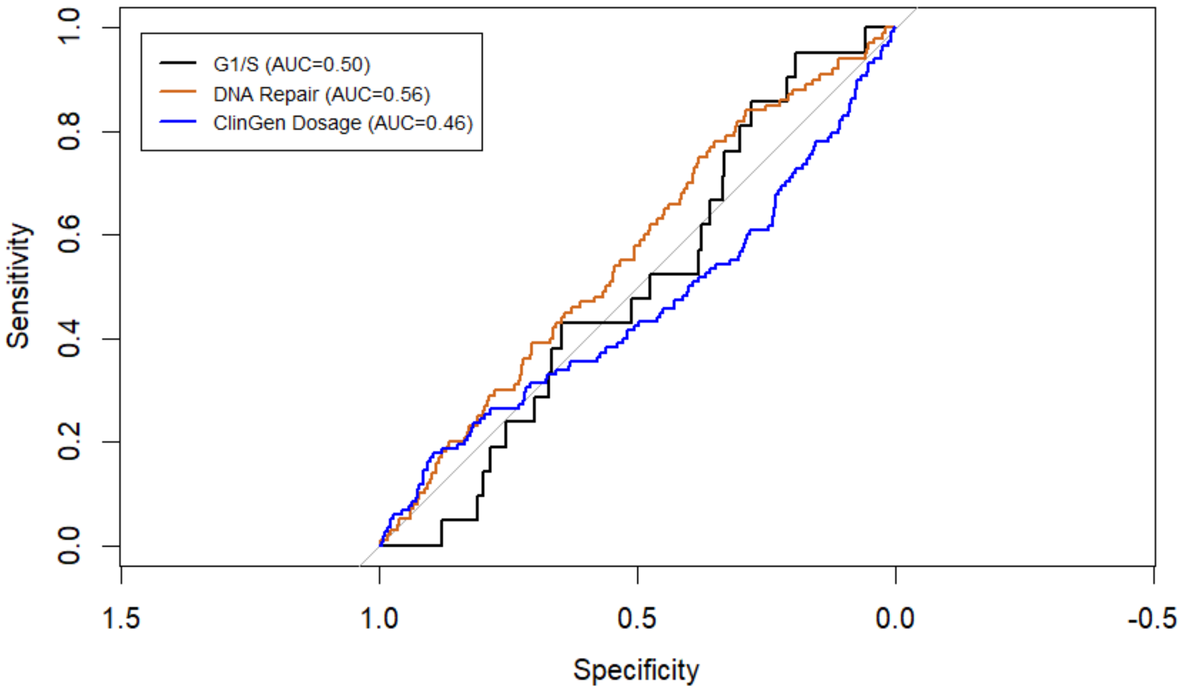


Supplementary Figure 10 | ROC Curve with OP->NO Intolerance (z-score) for multiple gene sets. Restricting to optimal-to-non-optimal changes modestly improved performance for DNA repair genes, but not for ClinGen or G1/S sets.

**10. Deriving the Relationship Between LRT Statistic and KL Divergence**

To interpret gene-level intolerance in a power-independent manner, we analytically derived the relationship between the multinomial likelihood ratio test (LRT) statistic used in CATMINT and the Kullback–Leibler (KL) divergence. Let $\theta_0$ and $\theta_1$ denote the expected and observed variant category proportions, respectively. Under the multinomial model, the LRT statistic comparing the null and alternative hypotheses is:

$$\Lambda = -2\ln\left(\frac{L(\theta_0)}{L(\theta_1)}\right)$$

Expanding this expression, we obtain:

$$\Lambda = -2ln\left(\frac{\left(\frac{n!}{x_1!\,x_2!\,x_3!\dots x_k!}\theta_{01}^{x_1}\theta_{02}^{x_2}\theta_{03}^{x_3}\dots\theta_{0k}^{x_k}\right)}{\left(\frac{n!}{x_1!\,x_2!\,x_3!\dots x_k!}\theta_{11}^{x_1}\theta_{12}^{x_2}\theta_{13}^{x_3}\dots\theta_{1k}^{x_k}\right)}\right)$$
$$= -2ln\left((\theta_{01}^{x_1}\theta_{02}^{x_2}\theta_{03}^{x_3}\dots\theta_{0k}^{x_k})/(\theta_{11}^{x_1}\theta_{12}^{x_2}\theta_{13}^{x_3}\dots\theta_{1k}^{x_k})\right)$$
$$= 2x_1 ln\left(\frac{\theta_{11}}{\theta_{01}}\right) + 2x_2 ln\left(\frac{\theta_{12}}{\theta_{02}}\right) + \dots + 2x_k ln\left(\frac{\theta_{1k}}{\theta_{0k}}\right)$$

Because of multinomial distribution, $x_i = n\theta_{1i}$, we obtain:

$$\Lambda = 2n\theta_{11}\, ln\left(\frac{\theta_{11}}{\theta_{01}}\right) + 2n\theta_{12}\, ln\left(\frac{\theta_{12}}{\theta_{02}}\right) + \dots + 2n\theta_{1k}\, ln\left(\frac{\theta_{1k}}{\theta_{0k}}\right)$$

$$= 2n\sum_i \theta_{1i}\ln\left(\frac{\theta_{0i}}{\theta_{1i}}\right) = 2n\, D_{KL}(\theta_1||\theta_0)$$

This derivation shows that LRT statistic is mathematically equivalent to the KL divergence between observed and expected variant distributions, scaled by the total variant count (also see Supplementary Figure 11). Because KL divergence captures the deviation in variant composition without being directly affected by sample size, we use it as a power-independent measure of intolerance throughout our analysis.

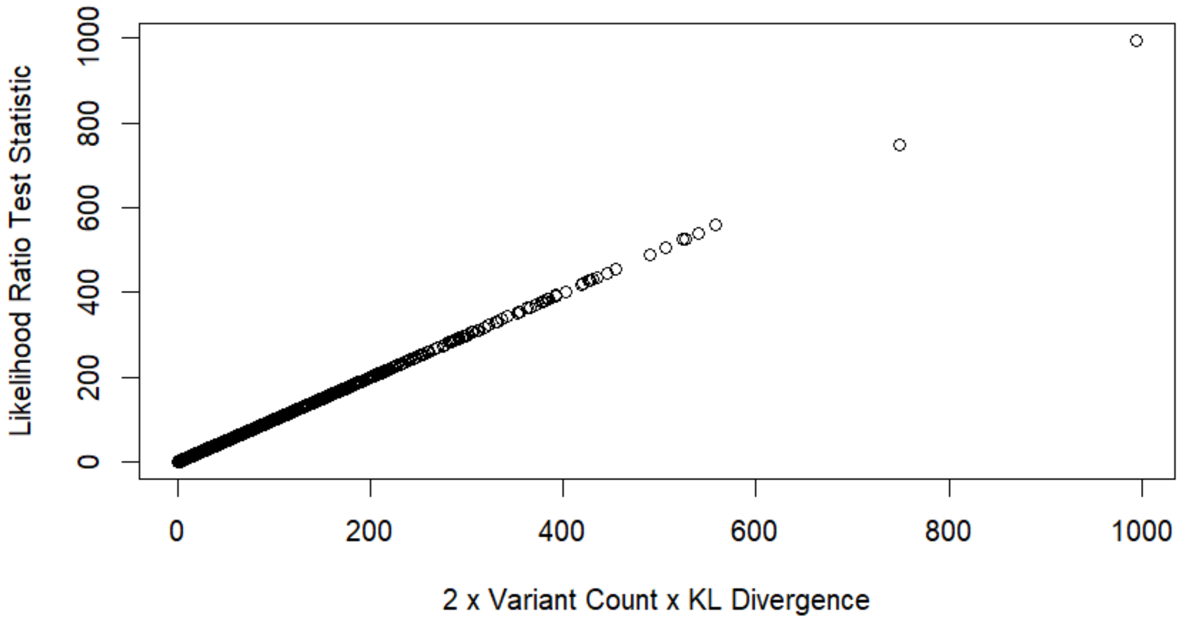


Supplementary Figure 11 | Empirical relationship between the CATMINT likelihood ratio test statistic and KL divergence. The LRT statistic from the CATMINT model (Y) aligns closely with 2 × variant count × KL divergence (X), confirming the analytical equivalence.

### 11. KL Divergence Thresholds Selection

To interpret variant composition deviation from neutrality independently of power, we selected thresholds for KL divergence based on differences in distribution between essential gene sets and all genes. We examined the cumulative distribution functions (CDFs) of KL divergence for several gene sets (Supplementary Figure 11). The intolerant gene sets, including stillbirth and haploinsufficient genes, showed consistent shifts toward higher KL divergence compared to all genes, suggesting stronger deviation from expected variant compositions. However, olfactory receptor genes showed less deviation.

To identify thresholds that distinguish these gene sets from the background, we calculated the enrichment of genes with more extreme KL divergence by subtracting the CDF of all genes from that of two representative essential gene sets: stillbirth genes and OMIM haploinsufficient genes (Supplementary Figures 12 and 13). Based on these enrichment curves, we selected a lower threshold of 0.005, representing the lower end of deviation observed among essential genes, also right before the CDFs of essential and all genes show large divergence. A higher threshold of 0.03 was selected to represent substantial deviation, based on a level commonly reached by the essential gene sets. These thresholds allowed us to capture both mild and strong compositional deviations: for instance, $KL < 0.005$ includes only 6.1% of stillbirth genes and 17.4% of haploinsufficient genes (vs. 43.5% of all genes), while $> 0.03$ includes 42.4% and 41.1% of those gene sets respectively (vs. 12.2% of all genes). To ensure that these patterns were not due to chance, we repeated the same analysis on 100 randomly sampled gene sets of the same size, using stillbirth genes as example (Supplementary Figure 14). The enrichment observed in essential genes was consistently stronger than in the random sets. These thresholds were used to help classify genes by their level of deviation, enabling us to identify genes with strong deviation but limited variant counts, or the opposite.

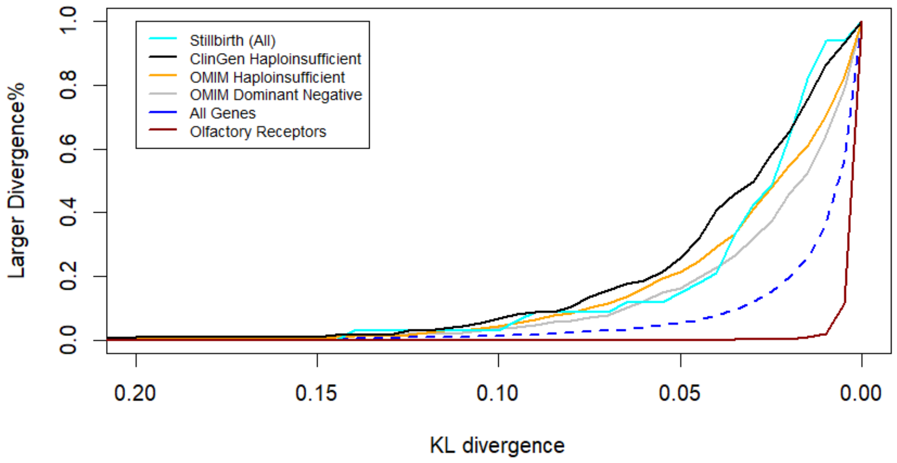


Supplementary Figure 12 | Proportion of genes with more extreme KL divergence (CDF of KL) in multiple gene sets. Stillbirth genes, ClinGen haploinsufficient genes, OMIM haploinsufficient genes and OMIM dominant negative genes showed more variant composition deviation from neutrality, compared with all genes, while olfactory receptor genes, which are known to be tolerant to variants, showed less deviation.

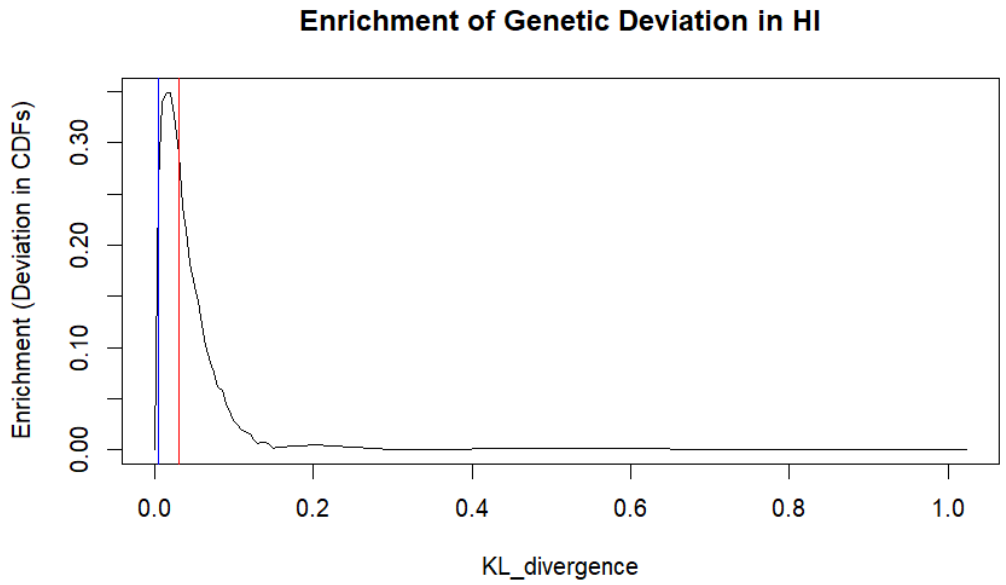


Supplementary Figure 13 | Enrichment of genetic deviation in haploinsufficient genes. KL divergence thresholds of 0.005 and 0.03 (blue and red lines) capture the points where enrichment deviates most from the background.

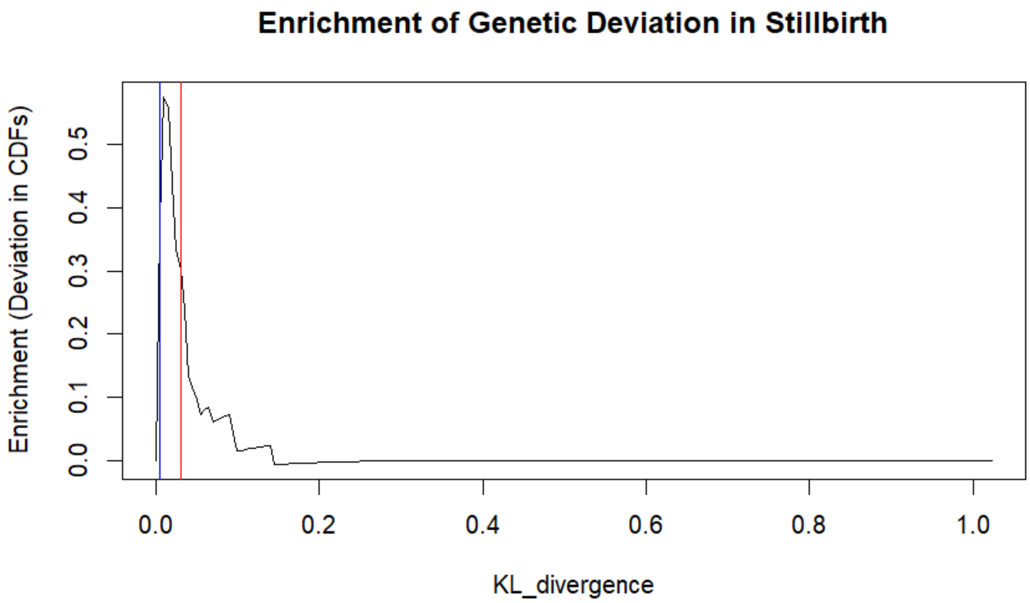


Supplementary Figure 14 | Enrichment of genetic deviation in stillbirth genes. Stillbirth genes also show strong enrichment between the 0.005 and 0.03 thresholds, similar to haploinsufficient genes.

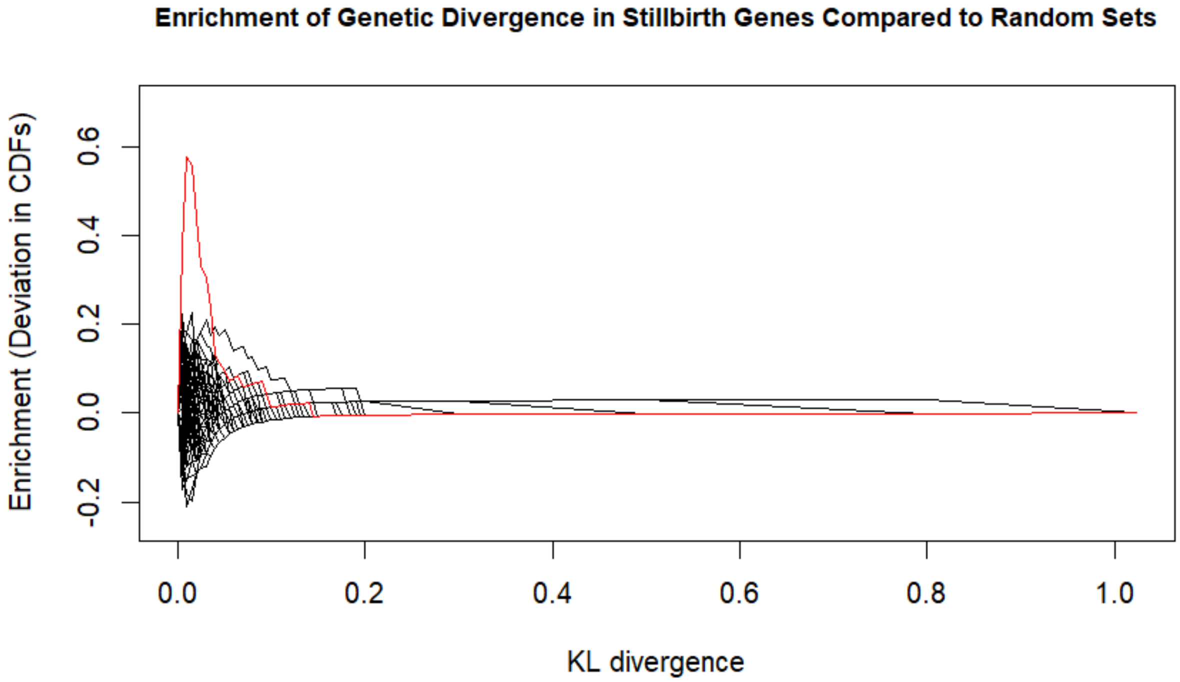


Supplementary Figure 15 | Enrichment of Genetic Divergence in Stillbirth Genes Compared to Random Sets. Stillbirth genes show greater enrichment than any random set, indicating the observed deviation is not due to chance.

**12. Enrichment of Essential Genes Among Small Genes**

We explored whether essential gene sets are biased toward longer genes and assessed how this bias affects the detection of intolerance in small genes. Gene length distributions across essential gene sets showed that ClinGen haploinsufficient and stillbirth genes tend to be longer, whereas MGI essential genes include a higher fraction of shorter genes (Supplementary Figure 16). Consistent with this, gene length and observed variant counts were highly correlated ($R^2 = 0.95$; Supplementary Figure 17), indicating that shorter genes naturally harbor fewer variants and may have lower statistical power to reach significance in CATMINT.

To evaluate whether small genes with strong deviation from expected variant composition are underrepresented in essential gene sets, we identified genes with KL divergence >0.03 but low variant counts (<301, >10) or short gene length (<519 bp). Enrichment analyses using Fisher's exact test showed that some essential gene sets, such as OMIM Recessive and neurodevelopmental genes, are depleted for these small but highly deviated genes, while MGI essential genes show mild enrichment (Supplementary Tables 7-8; Supplementary Figures 18-19). These patterns likely reflect differences in how the gene sets were constructed: literature-curated sets are more sensitive to detection biases, whereas experimentally derived sets (e.g., MGI essential) are less constrained by gene size.

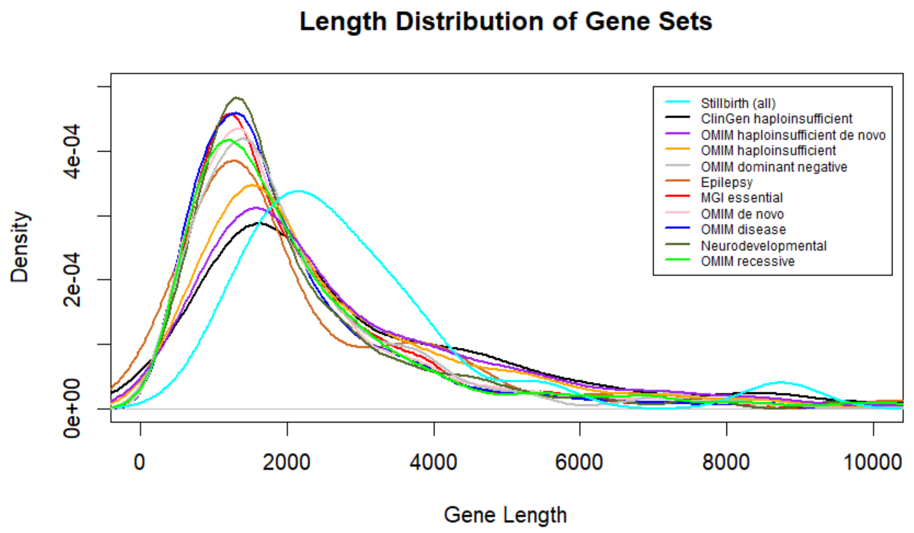


Supplementary Figure 16 | Gene Length Distribution in Essential Gene Sets. Density distributions of gene lengths for essential gene sets. Stillbirth and haploinsufficient genes show the strongest skew toward longer genes, whereas MGI essential genes are among the least biased toward long genes.

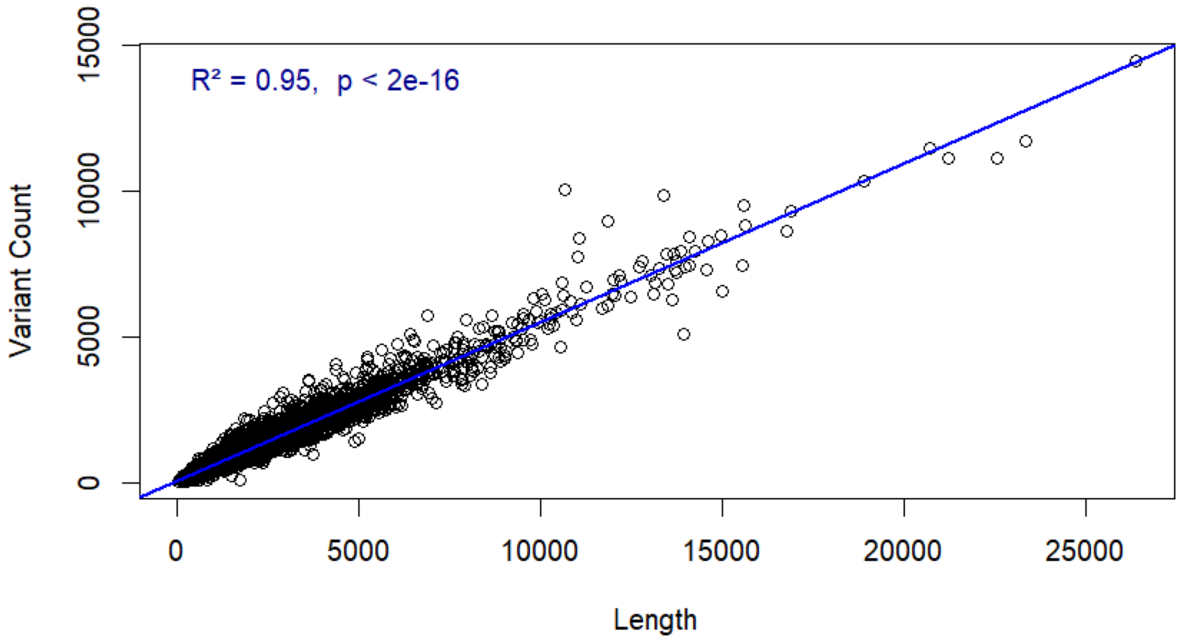


Supplementary Figure 17 | Correlation Between Gene Length and Number of Observed Variants (of high confidence genes). Scatter plot of observed variant counts versus gene length for all analyzed genes. A strong positive correlation was observed ($R^2 = 0.95$, $p < 2e-16$), justifying the use of gene length as a proxy for expected variant counts in defining “small” genes for enrichment analyses.

Supplementary Table 7 | Enrichment of small (by variant count), highly deviated genes in essential gene sets. Odds ratios and Fisher's exact test p-values for essential gene sets, evaluating enrichment among genes with variant count <301 (and >10) and KL divergence >0.03. These genes likely reflect underpowered but potentially intolerant genes.

| Gene Set | OR | p-value |
|---|---|---|
| MGI Essential Genes | 1.39 | 0.144 |
| Neuro Developmental Genes | 0.539 | 0.332 |
| OMIM Haploinsufficient Genes | 1.10 | 0.808 |
| OMIM De Novo Genes | 1.14 | 0.551 |
| OMIM HI De Novo Genes | 0.849 | 1 |
| OMIM Dominant Negative Genes | 1.17 | 0.673 |
| OMIM Recessive Genes | 0.25 | 0.00712 |

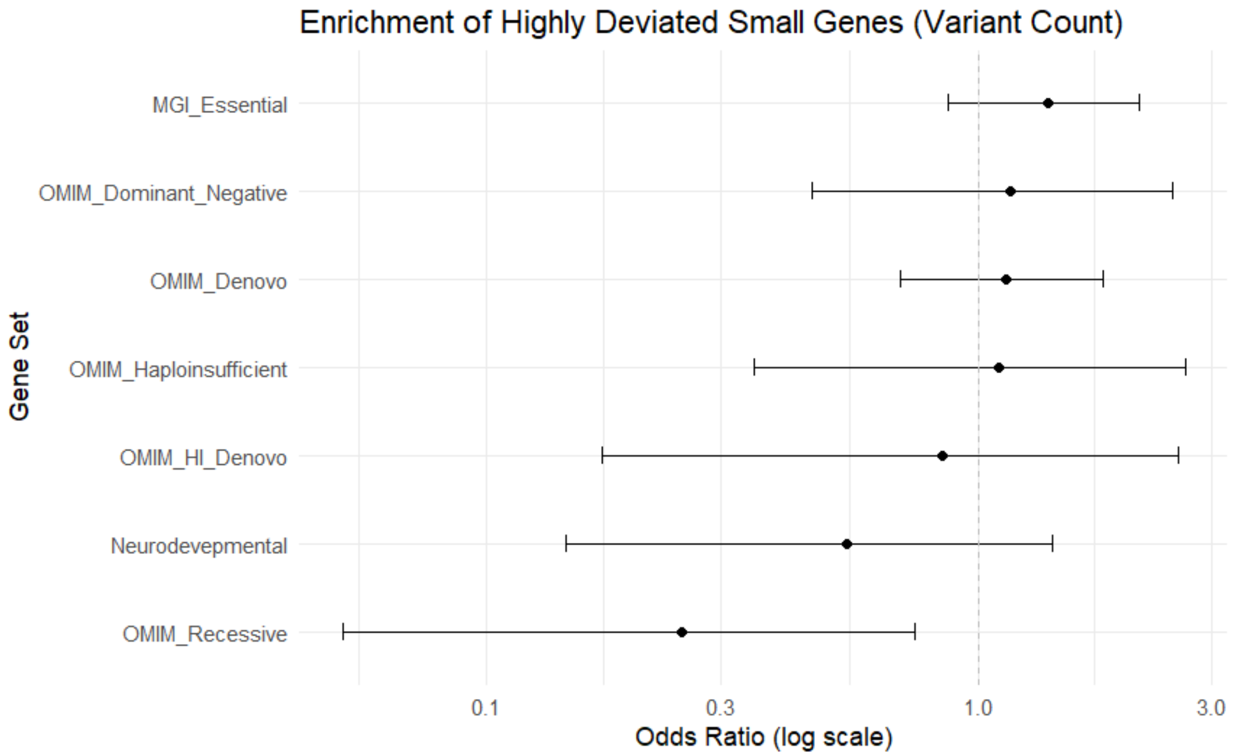


Supplementary Figure 18 | Enrichment of small (by variant count) , highly deviated genes in essential gene sets. Odds ratios (log scale) of essential gene sets for enrichment among genes with low variant counts (<301, >10) and high KL divergence (>0.03). MGI essential genes show slight enrichment, whereas OMIM recessive and neurodevelopmental genes are more depleted.

Supplementary Table 8 | Enrichment of small (by gene length), highly deviated genes in essential gene sets. Odds ratios and Fisher's exact test p-values for essential gene sets, evaluating enrichment among genes with short gene length (<519 bp, >10 variants) and KL divergence >0.03. This length-based definition identifies genes likely underpowered due to short length but with strong deviation from expected variant composition.

| Gene Set | EnF | p-value |
|---|---|---|
| MGI Essential Genes | 1.38 | 0.265 |
| Neuro Developmental Genes | 0.705 | 0.8 |
| OMIM Haploinsufficient Genes | 1.14 | 0.749 |
| OMIM De Novo Genes | 0.911 | 0.877 |
| OMIM Dominant Negative Genes | 0.848 | 1 |
| OMIM Recessive Genes | 0.439 | 0.217 |

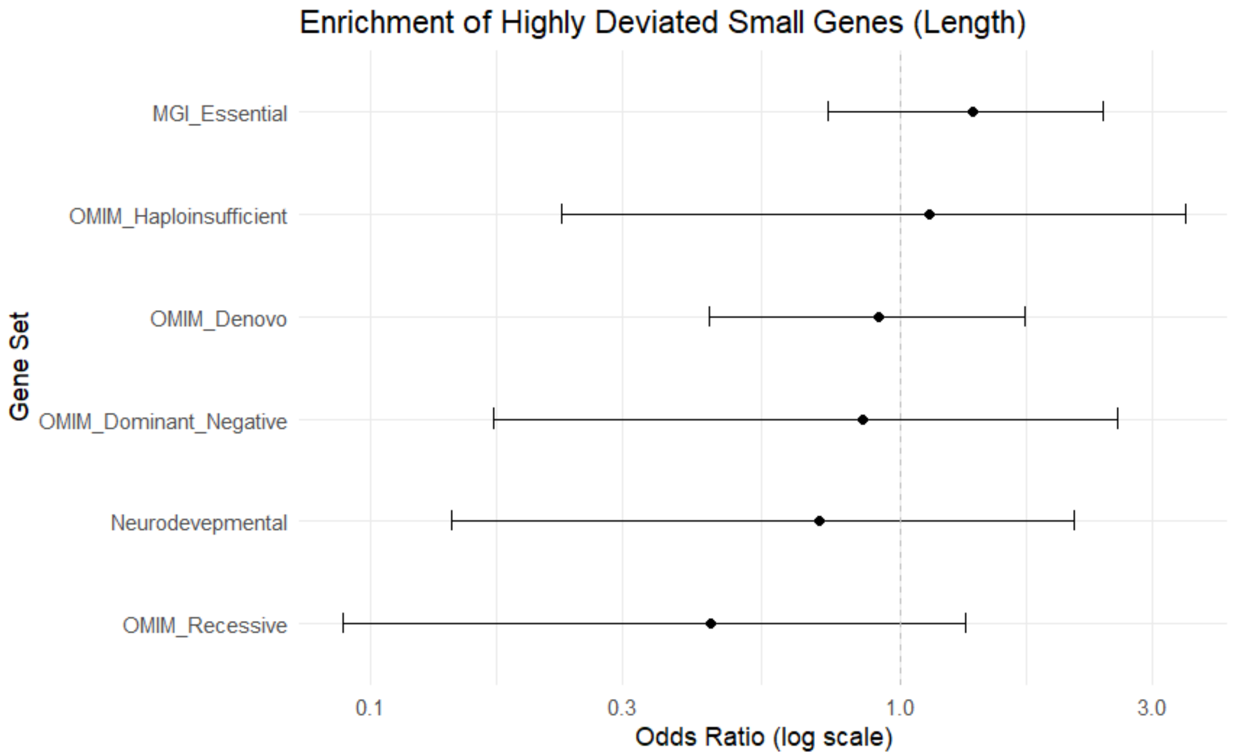


Supplementary Figure 19 | Enrichment of small (by gene length), highly deviated genes in essential gene sets. Odds ratios (log scale) showing enrichment or depletion of essential gene sets among short genes (<519 bp) with high KL divergence (>0.03). Patterns are consistent with variant-count–based analysis (Supplementary Figure 17), highlighting how gene size affects detection of intolerance.

### 13. Filtering Sequences for Balancing and Positive Selection Analysis

Whole genome sequencing (WGS) data were collected from 1000 Genomes Project. There were 3,202 samples consisting of individuals from 5 super-populations: African (AFR), Admixed American (AMR), European (EUR), East Asian (EAS) and South Asian (SAS) populations. Parents information was derived from a pedigree file at IGSR data portal. After removing parental IDs, 2,003 individual samples including 486 AFR, 220 AMR, 419 EUR, 443 EAS and 435 SAS remained for further analysis. Four kinds of repeats, low complexity repeats (LCR), Blacklist, centromere, and telomere were removed during the preprocessing steps (Data Availability).
